# Single-cell spatial map of cis-regulatory elements for disease-related genes in the macaque cortex

**DOI:** 10.1101/2025.06.03.657577

**Authors:** Juan Meng, Cheng Chen, Zhiyong Zhu, Yongkang Sun, Yiming Huang, Kaijie Hu, Jiqiang Fu, Luyan Wu, Ling Li, Yiqin Bai, Tianyi Fei, Zhen Liu, Chao Li, Zhiming Shen, Longqi Liu, Chengyu Li, Tao Song, Cirong Liu, Muming Poo, Shiping Liu, Ying Lei, Yidi Sun

## Abstract

Single-cell spatial transcriptomes have demonstrated molecular and cellular diversity in the brain^1, 2, 3, 4, 5, 6, 7, 8^, but gene regulatory mechanisms underlying transcriptomic profiles and disease pathogenesis remain largely unknown in primates. Here we performed single-nucleus Assay for Transposase-Accessible Chromatin followed by sequencing (snATAC-seq) for ∼1.6 million cells from 142 cortical regions of two male cynomolgus monkeys (Macaca fascicularis), and identified distinct chromatin accessibility profiles of cis-regulatory elements (CREs) for various cell types. By integrative analysis with large-scale spatial transcriptome data, we found that these CREs showed laminar and regional preferences, with their regional accessibility exhibiting striking dependence on the region’s hierarchical level. Cross-species comparison of snATAC-seq data revealed human/macaque-enriched layer-4 glutamatergic neurons and LAMP5/LHX6-expressing GABAergic neurons as well as human/macaque-biased CREs for genes related to neurodevelopment and psychiatric diseases. Importantly, risk single-nucleotide polymorphisms for many brain disorders strongly associated with human/macaque-biased CREs in glutamatergic neuronal types and those for Alzhemer’s disease strongly associated with CREs exclusively in microglia. Our results provided the basis for understanding the spatial gene regulatory mechanisms underlying cellular diversity and disease pathogenesis in the primate cortex.

## Introduction

Single-cell transcriptome analyses have facilitated our understanding of the diversity of cell types and their spatial distributions in the brain. Hundreds of glutamatergic, GABAergic, and glial cell types have now been identified^1, 2, 3, 4, 5, 6, 7, 8^. However, the gene regulatory mechanisms underlying their identity and function remain largely unclear. The spatiotemporal pattern of gene expression depends on the chromatin accessibility of cis-regulatory elements (CREs), which are binding sites for sequence-specific transcription factors (TFs) that recruit proteins for transcriptional initiation of their target genes. A comprehensive characterization of CREs at the single-cell level will help to elucidate gene regulatory mechanisms in diverse cell types and the molecular basis of brain development, function and diseases.

Recent studies using the snATAC-seq method have enabled the analysis of cell type-specific CREs in the adult mouse brain^9, 10, 11, 12^. Previous bulk sequencing assays have revealed that enhancer elements, the main component of CREs, exhibit rapid changes during mammalian evolution^13^ and primate-specific enhancer elements were found to be involved in brain diseases^14^. Thus, new CREs could appear during primate evolution and specifically responsible for human diseases. Indeed, comparative studies of single-cell chromatin accessibility datasets have shown conserved chromatin accessibility and CREs in human, monkey and mouse brains^15, 16^. However, whether primate-specific CREs underlie transcriptomic profiles of primate-enriched cell types and human disease pathogenesis remains largely unknown. Moreover, chromatin accessibility in bulk tissues was found to exhibit region-selective patterns in the primate brain^17^ and single-cell chromatin accessibility has been examined for a limited number of primate brain regions^15, 18, 19^. Thus, a comprehensive characterization of single-cell chromatin accessibility covering the entire macaque cortex will help to advance our understanding of CREs underlying the diversity of cell types and their regional distribution in the primate brain.

In this study, we generated a single-nucleus chromatin accessibility dataset for ∼1.6 million single nuclei from 142 cortical regions of the adult macaque brain using the snATAC-seq method. We defined 230 clusters of cells based on this chromatin accessibility atlas and identified 615,873 candidate CREs associated with various cell types in the macaque brain. Mapping of CREs with the spatial transcriptome showed their laminar and regional preferences. Further cross-species comparison of single-cell chromatin accessibility data from macaques, mice, and humans revealed human/macaque-biased CREs for psychiatric disorder-associated genes in layer-4 (L4) glutamatergic cell types and for neurogenesis-related genes in GABAergic neurons. Notably, we also found a strong association between human/macaque-biased CREs in specific cell types and risk single nucleotide polymorphisms (SNPs) for various brain diseases. These results provide a comprehensive database for studying gene regulatory programs at the single-cell level and for understanding pathogenic mechanisms underlying brain disorders. An interactive website for accessing our dataset is available through https://macaque.digital-brain.cn/chromatin-accessibility.

## Results

### Characterization of chromatin accessibility in various macaque cortical cell types

#### Processing of snATAC-seq data

To understand gene regulatory programs underlying cellular diversity in the primate cortex, we performed chromatin accessibility profiling using a previously reported droplet-based snATAC-seq method^20^ for tissue samples from 142 cortical regions (128 from Macaque #1 and 132 from Macaque #2) covering the entire cortex of two adult male cynomolgus monkeys (*Macaca fascicularis*) (Fig. 1a and Supplementary Table 1). The data reliability was confirmed by the high numbers of uniquely mapped reads, high proportion of properly paired reads, high numbers of unique fragments, low doublet ratio, and consistent quality metrics between biological replicates (Extended Data Fig. 1a-e). A total of 1,615,430 nuclei passed quality control criteria including the number of unique fragments per cell, TSS enrichment score, promoter ratio and doublet removal (Methods and Extended Data Fig. 1f, g). The remaining cells exhibited an average of 20,534 chromatin fragments in each nucleus and 57.6% of reads in peak regions. These data were used for unsupervised clustering analysis, and assisted by previously reported spatial transcriptome dataset^21^, our chromatin accessibility data were used for further spatial mapping of CREs and gene regulatory networks (GRNs; Fig. 1a).

**Fig. 1.**
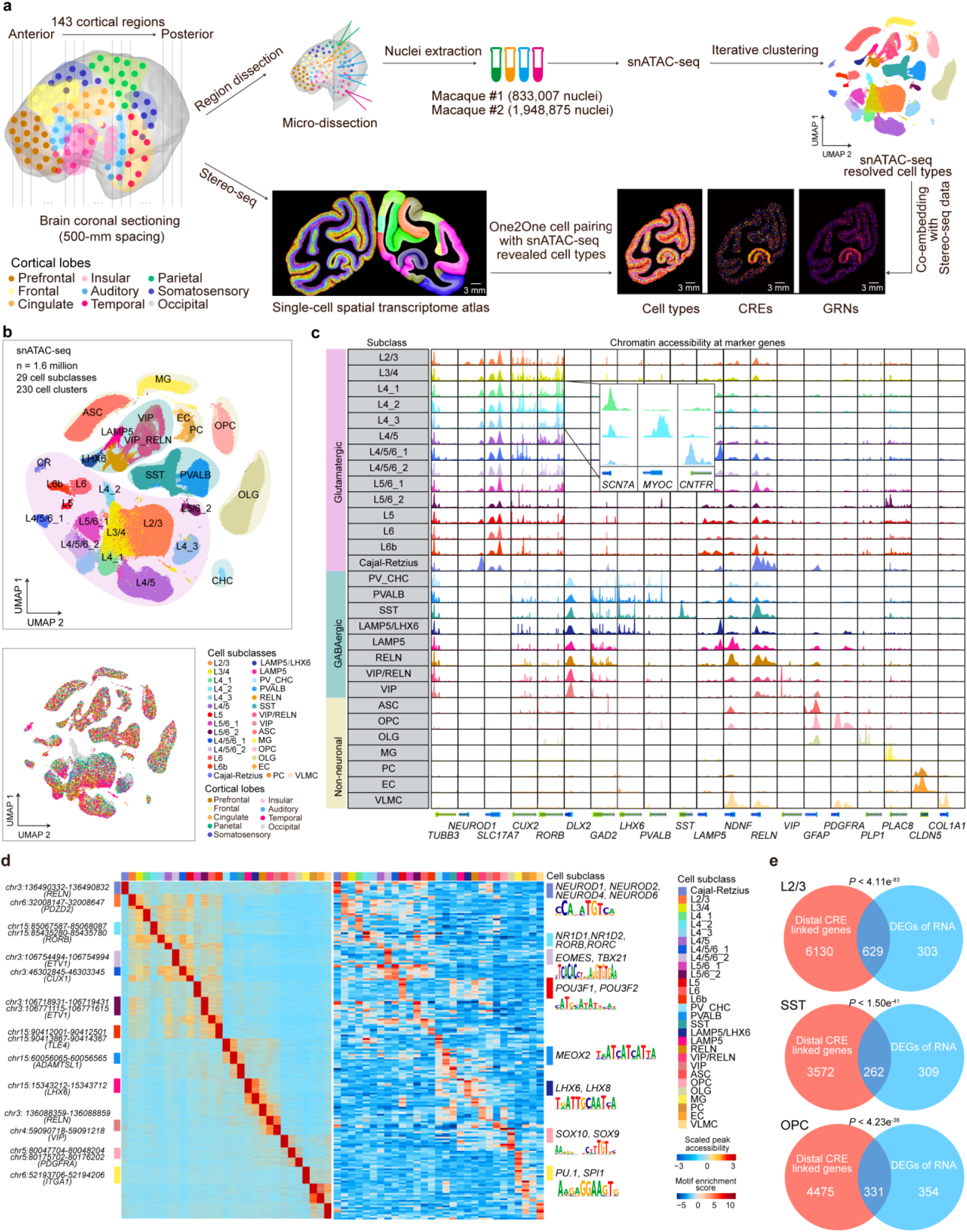
Chromatin accessibility profiles of various cell types in the macaque cortex. **a**, Schematics of experimental procedure and integration of snATAC-seq with snRNA-seq and spatial transcriptome data. Single nuclei from 142 cortical regions of two monkeys were isolated for snATAC-seq and snRNA-seq analysis, and co-embedded with spatial transcriptome data from a previous study^21^. **b**, Upper, UMAP embedding and clustering analysis based on chromatin accessibility profiles of snATAC-seq data. Below, UMAP layout showing single nuclei from different cortical lobes. Single nuclei are colored by cell clusters or cortical lobes. A full list and description of cell clusters and subclusters are provided in Supplementary Table 2. L2/3, layer 2/3, CR, Cajal-Retzius, PV, Parvalbumin, CHC, chandelier cell, SST, Somatostatin, ASC, astrocyte, OPC, oligodendrocyte precursor cell, OLG, oligodendrocyte, MG, microglia, EC, endothelial cells, PC, pericytes, VLMC, vascular leptomeningeal cells. **c**, Marker peaks of aggregated chromatin accessibility profiles for each snATAC-seq-based cell cluster at selected marker gene loci that were used for cell annotation. **d**, Left, Heatmap showing the chromatin accessibility of CREs across various cell clusters. Right, Heatmap showing the enriched TF motifs for various cell clusters. Representative TFs and their corresponding motif logos were shown on the right. **e**, Venn plots showing the overlap between distal CRE linked genes and differentially expressed genes in the same cell type. *P* values were calculated by hypergeometric test.

#### Clustering analysis of chromatin accessibility profiles

Unsupervised clustering based on the representations after iterative latent semantic indexing (LSI) of chromatin accessibility in each cell resulted in three major types, glutamatergic (726,611; 45%), GABAergic (275,073; 17%) and non-neuronal (613,746; 38%) cells. Further iterative clustering of each major type revealed 91 subclusters (14 clusters) of glutamatergic neurons, 105 subclusters (8 clusters) of GABAergic neurons, and 34 subclusters (7 clusters) of non-neuronal cells (Fig. 1b and Supplementary Table 2). The major clusters and subclusters were annotated by the peak accessibility at the promoter region of their canonical marker genes^6, 22, 23^. For examples, high chromatin accessibility was observed at the promoter region of *SLC17A7* for all glutamatergic neurons and *GAD2* for GABAergic neurons. Clusters associated with astrocytes (ASCs), oligodendrocyte progenitor cells (OPCs), oligodendrocytes (OLGs) and microglia cells (MGs) showed open chromatin peaks at the promoter of *GFAP*, *PDGFRA*, *PLP1* and *PLAC8*, respectively (Fig. 1c). Cells from multiple cortical areas of two monkeys were present in the majority of cell types (Fig. 1b), and high consistency in the proportions of various clusters was found between biological replicates (Extended Data Fig. 1h-k), demonstrating the reliability of the batch correction. Hierarchical clustering of the averaged LSI representations in each cluster resulted in a taxonomy tree (Extended Data Fig. 1l). Integrative analysis of snATAC-seq and snRNA-seq data showed that the clusters of chromatin accessibility profiles identified by snATAC-seq corresponded well with snRNA-seq-defined cell types (Extended Data Fig. 2a, b and Supplementary Table 2).

#### Identification of open chromatin regions in each snATAC-seq clusters

We further applied the MACS2 method^24^ to identify open chromatin regions for each cell type by aggregating chromatin accessibility profiles (Methods). After filtering out those with low confidence, we detected an average of 138,477 open chromatin regions per cell type and a total of 615,873 open chromatin regions in all cell types (Extended Data Fig. 2c). Among these open chromatin regions, more than 90% were located ±1 kb away from known transcriptional start site (TSS) of protein-coding and long noncoding RNA genes, consistent with previous data for human and mouse brains^11, 15, 18, 19^ (Extended Data Fig. 2d). Interestingly, with comparable numbers of chromatin fragments among different cell types, the proportions of open chromatin peaks enriched in the TSS or promoter regions (more adjacent cis-regions) in non-neuronal cells and GABAergic cells were much higher than those in glutamatergic cells (Extended Data Fig. 2e). Our analysis of previously published datasets of human, macaques and mice^11, 15, 18, 19^ showed the same phenomenon (Extended Data Fig. 2e). The lower TSS enrichment in glutamatergic neurons might be induced by higher activity of glutamatergic neurons or more distant regulatory elements from the TSS in glutamatergic cells. As a previous study^25^ has shown that neuronal activity induces a major reconfiguration of the chromatin state, with most of the activity induced open regions located in the intergenic and intronic regions.

#### Identification of CREs and their linked genes in various cell types

We thus defined these distal open chromatin regions as cis-regulatory elements (CREs) and linked them with putative target genes by calculating the co-accessibility of each distal CRE and open regions within ±1 kb from the TSS of targeted genes in each cell type using the Cicero method^26^, resulting in 1,872,039 CRE-gene pairs (Supplementary Table 3). Over 70% of CREs (436,615) showed differential chromatin accessibility across different cell subclasses, and a large proportion of CRE-linked genes in each cell subclass overlapped with marker genes in the corresponding snRNA-seq-defined cell subclasses (Fig. 1e, Extended Data Fig. 2f and Supplementary Table 3). For examples, L2/3 glutamatergic cell subclass showed high chromatin accessibility in CREs linked to *GPR83* and *PDZD2* genes, which were also highly expressed in snRNA-seq-defined L2/3 neurons. Moreover, these cell subclass-restricted CREs were also enriched for distinct sets of DNA motifs recognized by known transcriptional regulators (Fig. 1d). For instances, the motif binding activity of the nuclear receptor gene *NR1D2,* which encodes the TF involved in several brain disorders, showed the highest level in L4_2 glutamatergic neurons. The activity of binding motifs by SOX family TFs *SOX9* and *SOX10*, which are involved in oligodendrocyte specification and myelination^27^, were found to be highest in OPCs (Fig. 1d). These findings establish a basis for understanding gene regulatory mechanisms underlying various transcriptome-defined cell types in the macaque cortex.

### Spatial mapping of chromatin accessibility revealed by snATAC-seq data

#### Mapping chromatin accessibility of CREs to single cells on the spatial transcriptome map

To further examine the spatial accessibility of cell type-specific CREs, we integrated the snATAC-seq data with single-cell spatial transcriptome map of the macaque cortex^21^ for one-to-one pairing. Specifically, we down-sampled the “gene activity score” matrix of snATAC-seq data to mimic the gene capture rate of Stereo-seq data, and then used autoencoder and generative adversarial network to transfer snATAC-seq-identified clusters onto the spatial map (Fig. 2a). The optimal transport method was used at the same time to achieve one-to-one mapping, in which chromatin accessibility profiles from snATAC-seq data were assigned to single cells on the spatial transcriptome map (Methods). The accuracy of this cell mapping was evidenced by the high consistency in the relative percentage of different cell types and marker genes between the snATAC-seq and spatial transcriptome data for the same cell types (Extended Data Fig. 3a, b).

**Fig. 2.**
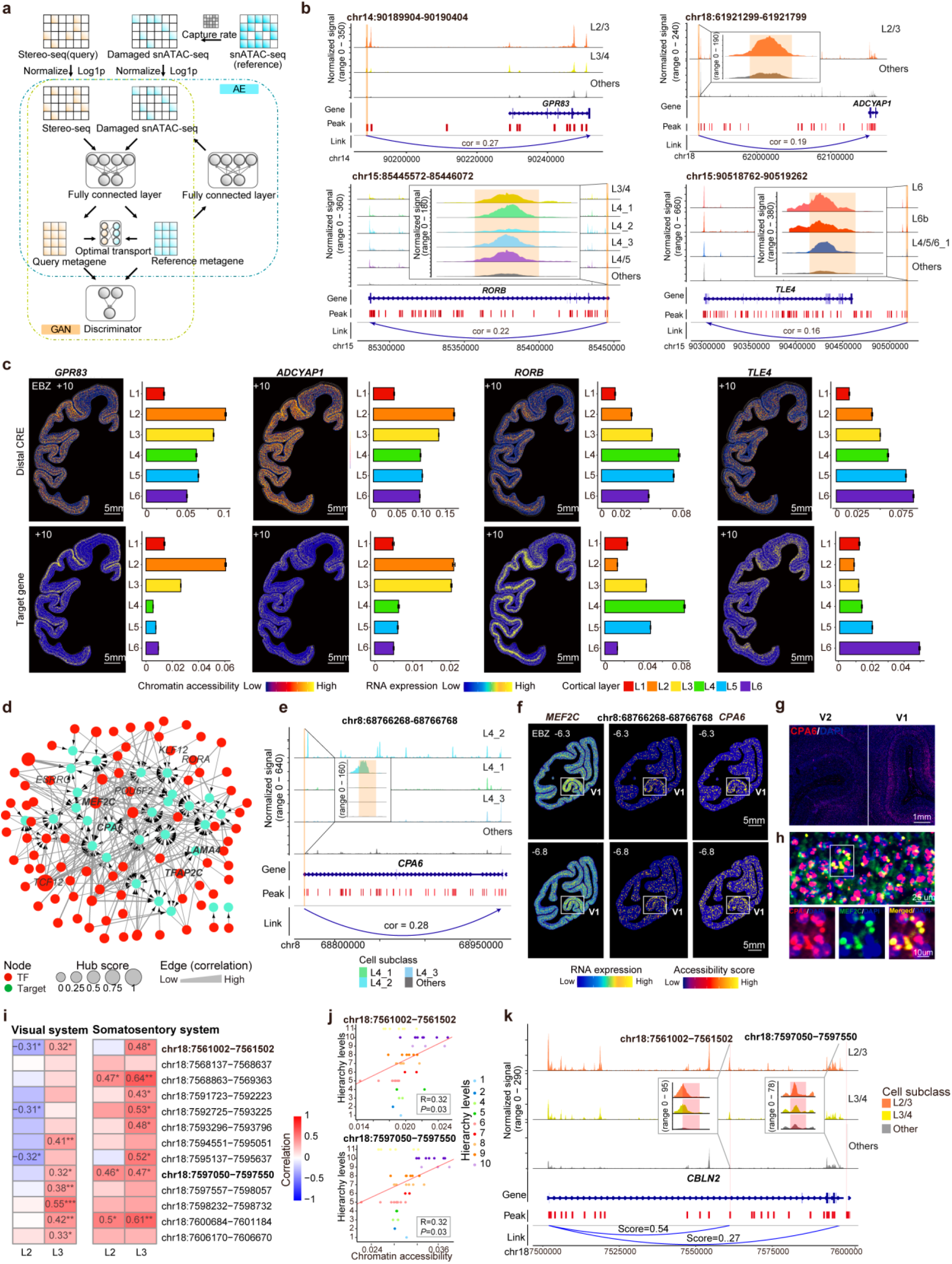
Characterization of CREs across various brain regions of the macaque cortex. **a**, Diagram showing the integration and one-to-one pairing of snATAC-seq and spatial transcriptome data. **b**, Genome browser track view showing differential accessible distal CREs linked to *GPR83*, *ADCYAP1*, *RORB* and *TLE4* in various glutamatergic neurons. **c**, Spatial maps and bar plots showing the chromatin accessibility of four example CREs and expression levels of these CREs-linked genes across cortical layers in a representative coronal section (EBZ +10). The quantification was calculated from chromatin accessibility in five representative coronal sections (Fig. 2c and Extended Data Fig. 3d). Error bars represent standard error of the mean (SEM). **d**, Gene regulatory networks of the V1-specific L4_2 glutamatergic cell cluster. Red and green colors represent TFs and their target genes, respectively. The directed edges indicate the regulation between TFs and their targeted genes. **e**, Genome browser track view showing a differential accessible distal CRE linked to *CPA6* across three L4 glutamatergic clusters and other glutamatergic clusters. **f**, Spatial maps of expression patterns of *MEF2C* and *CPA6*, and chromatin accessibility of the CRE linked to *CPA6* at two representative sections (EBZ -6.3 and -6.8). **g**, RNAscope assay showing the expression pattern of the *CPA6* gene in V1 and V2 regions of the macaque cortex. Probes against CPA6 (ACD, cat#1843691-C1) was list in Supplementary Table 8. DAPI was used for nuclear counterstaining. **h**, Co-localization of RNAscope labeling of the *CPA6* gene and the *MEF2C* gene in the V1 region. **i**, Heatmap showing the correlation of chromatin accessibility and the hierarchy levels of various brain regions in the visual and somatosensory system. Correlation coefficients and *P* values were calculated by Spearman correlation analysis. * *P* < 0.05, ** *P* < 0.01, *** *P* < 0.001. **j**, Dot plots showing the correlation of chromatin accessibility and the hierarchy levels of various brain regions in the visual system. **k**, Genome browser tracks showing the chromatin accessibility of two CREs linked to *CBLN2* among L2/3, L3/4 and other cell subclasses.

#### Cortical layer preference of chromatin accessibility of CREs

To explore the spatial distribution pattern of chromatin accessibility for cell type-specific CREs, we then calculated the relative percentage of each cell type in each of the six cortical layers and found that glutamatergic subclusters showed clear laminar preference (Extended Data Fig. 3c). The cell type-specific CREs also showed laminar-specific chromatin accessibility in the spatial map, consistent with the preferential laminar expression of their targeted genes. For example, the CRE linked to the marker gene *GPR83* of L2/3 glutamatergic neurons showed open chromatin accessibility peaks only in L2/3 (Fig. 2b). Spatial mapping of this CRE also showed significantly higher chromatin accessibility in L2 and L3, consistent with the spatial expression preference of *GPR83* (Fig. 2c and Extended Data Fig. 3d). In addition, distal CREs linked to *ADCYAP1*, *RORB*, and *TLE4* also showed consistent preferences in spatial chromatin accessibility with the expression pattern of corresponding marker genes for L2/3, L4 and L6 glutamatergic neurons, respectively (Fig. 2b, 2c and Extended Data Fig. 3d).

#### Cortical region preference of chromatin accessibility of CREs

In addition to layer-specific distribution, we also found cortical region-specific distribution of some cell clusters based on their relative density among nine cortical regions (Extended Data Fig. 3c). For example, the L4_2 glutamatergic subclass exhibiting high chromatin accessibility at the promoter of *MYOC* was enriched in the V1 region of the occipital lobe quantified from both snATAC-seq and spatial transcriptome data (Fig. 1c and Extended Data Fig. 3e, f). This cluster corresponded well to snRNA-seq-defined L4.4 and L4.8 cell types that were also specifically distributed in V1^21^ (Extended Data Fig. 3g). The DNA sequences of CREs enriched in L4_2 glutamatergic cluster were then scanned for TF-binding motifs with the CellOracle method^28^, generating a gene regulatory network (GRN) for all potential regulatory interactions (Fig. 2d). One marker CRE linked to the *CPA6* gene was significantly enriched with the binding motif for TF MEF2C (Fig. 2e), which is involved in normal neuronal development, cell distribution, and electrical activity of the neocortex as well as neuropsychiatric disorders^29^. Spatial mapping of this CRE also showed that its chromatin accessibility was V1-specific (Fig. 2f). Furthermore, the RNAscope analysis validated the V1-specific expression of *CPA6* and higher expression of *MEF2C* in V1 than V2 (Fig. 2g and Extended Data Fig. 3h). In addition, co-localization of the TF MEF2C and its target gene *CPA6* were found in the V1 region (Fig. 2h), suggesting their V1-specific regulation. Similarly, both the chromatin accessibility of CRE linked to *LAMA4* and the expression of *LAMA4* showed V1 specificity (Extended Data Fig. 3i, j). These results indicate that the laminar- and regional-specific accessibility of CREs might underlie the specific expression patterns of their targeted genes.

#### Characterization of CREs in the piriform cortex

As the evolutionarily ancient part of the cerebral cortex in the brain, the allocortex has simpler structure than the more common neocortex^30^. To understand the regulatory differences between the allocortex and neocortex, we next performed differential CRE analysis between the allocortical piriform cortex and all the other neocortical regions. A total of 19,376 CREs showed significantly differential chromatin accessibility between the two regions (Extended Data Fig. 4a). The up-regulated CREs in the piriform cortex were enriched in oligodendrocytes (OLGs) and other glial cells, whereas the down-regulated CREs were mainly contributed by deep-layer glutamatergic neurons (Extended Data Fig. 4b). Moreover, the piriform cortex included a higher proportion of glial cells (especially OLGs) and lacked L4 neuronal subtypes (L4_1, L4_2 and L4_3) (Extended Data Fig. 4c), in line with the anatomical absence of layer 4 in the piriform cortex. In addition, GO functional enrichment analysis revealed that genes linked by up-regulated CREs (*CSF1R*, *P2RY1*, *CX3CR1*) were significantly enriched in the “gliogenesis” term, whereas genes linked by down-regulated CREs (*KCNIP4*, *RGS7*, *SLC24A2*) were enriched in “potassium ion transport” term (Extended Data Fig. 4d, e). Spatial mapping of these pathways showed that the “gliogenesis” pathway was significantly enriched in the piriform cortex, whereas the “potassium ion transport” showed a lower abundance than other neocortical lobes (Extended Data Fig. 4f, g). As examples, the CRE linked to *CSF1R* showed high chromatin accessibility in the piriform cortex (Extended Data Fig. 4h). In contrast, the CREs linked to *KCNIP4* showed high chromatin accessibility in the other neocortex (Extended Data Fig. 4i). Therefore, these results revealed different regulatory programs between the piriform and neocortical regions.

#### Gradients of CRE accessibility across the neocortex

Given the laminar and regional preferences of CRE accessibility, we next wondered whether the accessibility of cell type-specific CREs showed intracortical gradients along the neocortex. As an example, the *CBLN2* is known to exhibit an anterior-posterior gradient in expression enriched in the prefrontal cortex during both development and adulthood in macaques and humans, and its enhancer has been identified to be active in the prefrontal cortex^31^. In order to assess the potential gradients of CREs associated with *CBLN2* along the macaque cortical hierarchy, we mapped the CREs linked to *CBLN2* gene in our spatial transcriptome map and found their higher chromatin accessibility in L2 and L3 (Extended Data Fig. 4j, k). We further performed Spearman correlation analysis of chromatin accessibility and the hierarchy levels of various brain regions in the visual system and the somatosensory system. The results showed that the accessibility of most of these CREs in L3 exhibited a significantly positive correlation with the hierarchy level of brain regions in both the visual system and the somatosensory system (Fig. 2i). Two example CREs (chr18:7561002-7561502 and chr18:7597050-7597550) showed open chromatin peaks in L2/3 and L3/4 and exhibited increasing accessibility along the hierarchy of both visual and somatosensory systems in L3 (Fig. 2j, k and Extended Data Fig. 4l). Taken together, these results revealed differences in CRE accessibility across cortical layers and regions, supporting the role of different regulatory programs in the spatial organization of various cell types in the primate neocortex.

### Human/macaque-biased CREs in glutamatergic neurons

#### Cross-species comparison of chromatin accessibility

To determine whether the gene regulation landscape in the cortex are conserved among humans, macaques and mice, we retrieved previously published snATAC-seq data for human and mouse cortices^11, 15^. Joint clustering analysis was performed using gene activity scores of homologous genes in snATAC-seq datasets for cortical tissues from all three species. We found generally conserved snATAC-seq-based clusters of GABAergic and non-neuronal cells across the three species, except for L4_1, L4_3 and L4/5/6_2 glutamatergic neurons that existed only in humans and macaques, as shown by the extracted maps of clusters for each species (Fig. 3a, Extended Data Fig. 5a-b and Supplementary Table 4). Further quantitative analysis among the three species using MetaNeighbor^32^ showed that macaque L4_1, L4_3 and L4/5/6_2 neurons showed high similarity with human L3/4 IT, L4 IT and L4/5 IT cell clusters, respectively, but no correspondence to any cluster in mice (Fig. 3b). Further comparison of all chromatin accessibility clusters revealed 12 clusters (belonging to L4_1, L4_3 and L4/5/6_2 subclasses) that were exclusively identified in primates but not in mice (Extended Data Fig. 5c). The human/macaque-biased clusters also corresponded well to snRNA-seq-based L4 and L4/5/6 glutamatergic cell types (Extended Data Fig. 3g), which were previously shown to be primate-specific^21^.

**Fig. 3.**
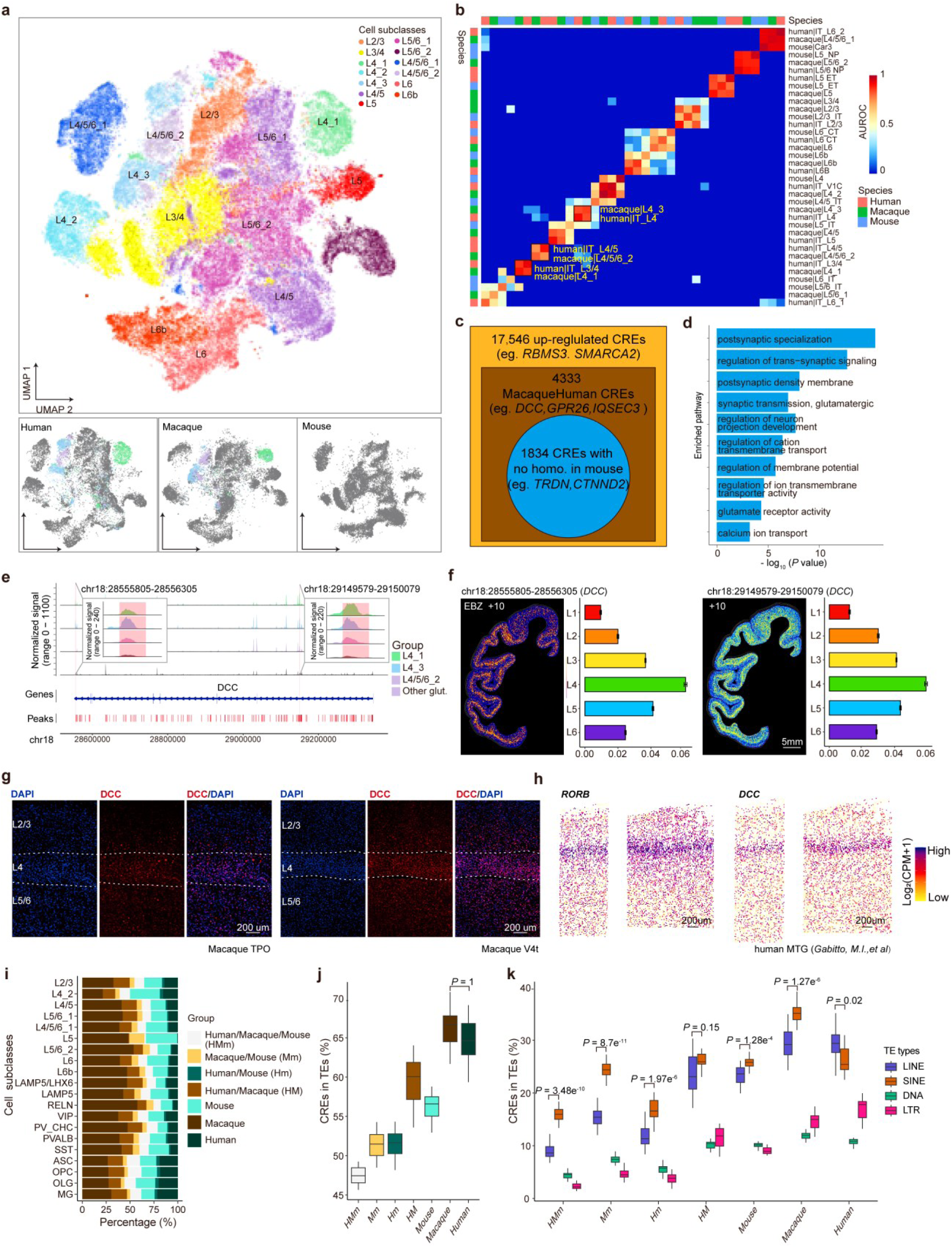
Comparative analyses revealed human/macaque-biased CREs in glutamatergic neurons. **a**, UMAP co-embedding of snATAC-seq data from macaques, humans and mice. Colored dots at the bottom indicate human/macaque-biased glutamatergic neurons derived from snATAC-seq data. **b**, Heatmap showing the similarity of snATAC-seq derived glutamatergic cell clusters across macaques, humans and mice using MetaNeighbor. **c**, The outer box represents the number of CREs showing higher chromatin accessibility in human/macaque-biased than other cell types of the macaque cortex. The inner box represents CREs with high chromatin accessibility in the corresponding human cell types. The inner circle indicates CREs showing no homologous sequence in the mouse genome. **d**, GO enrichment for genes linked to human/macaque-biased CREs (inner circle in **c**). **e**, Genome browser tracks showing the chromatin accessibility of human/macaque-biased CREs located on the *DCC* gene in the human/macaque-biased and other cell subclasses. **f**, Spatial map showing the chromatin accessibility of the human/macaque-biased CREs shown in **e** in a representative Stereo-seq section at indicated coordinate. The right panel showing the quantitative chromatin accessibility level across six cortical layers. Error bars represent standard error of the mean (SEM). **g,** RNAscope results showing the layer 4-specific expression of *DCC* gene in TPO (left) and V4t (right) regions of the macaque cortex. Probes against *DCC* (ACD, cat#1843701-C1) was listed in Supplementary Table 8. DAPI was used for nuclear counterstaining**. h,** Scatterplots showing the expression of layer 4 marker gene *RORB* and *DCC* gene in human MTG of the previously published MERFISH dataset^34^. **i**, Bar plots showing 7 categories of CREs in various cell subclasses from macaques, humans and mice. **j**, Boxplots showing the percentage of CREs colocalized with TE in different categories. **k**, Boxplots showing the percentage of CREs located in different types of TEs. *P* values were calculated by Wilcoxon rank-sum test.

#### Identification of human/macaque-biased CREs linked to disease associated genes

Further differential CRE analysis between human/macaque-biased cell types and all other glutamatergic cell types identified 17,546 CREs showing higher chromatin accessibility in the human/macaque-biased than other cell types (Extended Data Fig. 5d). Among these CREs, 4,333 CREs also showed high open chromatins in the corresponding human cell types, and 1,834 CREs had no homologous sequences in mice (Fig. 3c, Extended Data Fig. 5e and Supplementary Table 4). These human/macaque-biased CREs linked genes were significantly enriched in “glutamatergic synaptic transmission”, “postsynaptic specialization”, "regulation of trans-synaptic signaling" and “postsynaptic density membrane” (Fig. 3d). Specifically, we found that the chromatin accessibility of two CREs located in the intron of the neurodevelopmental disorder-associated gene *DCC* (netrin 1 receptor)^33^ were much higher in these human/macaque-biased neurons than those in other cell types (Fig. 3e), consistent with the higher expression level of *DCC* in L4, L3/4, L3/4/5 and L4/5 glutamatergic neurons (Extended Data Fig. 5f). These CREs showed much higher accessibility in L4 for most coronal regions and in L3 and L5 for regions without L4 quantified by spatial mapping (Fig. 3f and Extended Data Fig. 5g), consistent with the expression pattern of *DCC*. RNAscope assay verified that the expression of *DCC* was enriched in layer 4 (Fig. 3g). Analysis using the previously published MERFISH dataset from the human MTG region^34^ further validated the layer 4-specific expression of *DCC* gene, which is consistent with the expression pattern of *RORB*, a canonical marker gene of layer 4^34^ (Fig. 3h). Another example is a distal CRE that linked to *TRDN* with higher chromatin accessibility in these human/macaque-biased glutamatergic neurons (Extended Data Fig. 5h). The decreased expression of *TRDN* was reported to be associated with apoptosis of dopaminergic cells in Parkinson’s disease^35^. Spatial mapping showed the higher chromatin accessibility of this CRE and higher expression level of *TRDN* in deeper layers (Extended Data Fig. 5i, j). These results suggested the unique roles of human/macaque-biased CREs in regulating the spatial expression of disease-associated genes in the primate cortex.

Furthermore, we also examined whether CREs identified in macaque cell types that were conserved across three species were present in human or mouse cortices. For this analysis, we lifted CREs identified in each species to the human genome by sequence homology, and compared the sequence similarity and chromatin accessibility of CREs in each conserved cell subclass. Only CREs showing both sequence similarity and open chromatin between species were considered as conserved (Methods). We found that in comparison with species-conserved CREs, a higher proportion of CREs showed open chromatin in a species-specific pattern (Fig. 3i). Interestingly, we found that ∼60% of human/macaque-biased CREs were colocalized with known transposable elements (TEs) (Fig. 3j), consistent with the notion that TEs as a driver of genomic diversity^36^. In comparison with CREs conserved between primates and mice, human/macaque-biased or species-specific CREs showed significantly higher frequency in co-localization with TE elements including LINE, SINE, DNA and long terminal repeats. In addition, human/macaque-biased CREs showed comparable enrichment with LINE and SINE, whereas conserved CREs between primates and mice showed much higher co-localization with SINE element (Fig. 3k). Thus, we have identified human/macaque-biased CREs that may determine layer- and region-specific glutamatergic cell types and may be responsible for gene regulation underlying brain disorders.

### Characterization of CREs in GABAergic cell types

#### Identification of a human/macaque-enriched LAMP5/LHX6 GABAergic cell type

The integrative clustering analysis also showed clear correspondence of snATAC-seq identified clusters and snRNA-seq-defined GABAergic cell types (Extended Data Fig. 6a). However, based on chromatin accessibility profiles, we found that transcriptome-defined LAMP5+ GABA cell subclass could be further divided into LAMP5 and LAMP5/LHX6 cell types that exhibited low and high chromatin accessibility of the *LHX6* gene, respectively. We identified 3,829 and 1,989 differentially accessible (DA) chromatin peaks in LAMP5/LHX6 and LAMP5 cells, respectively. GO analysis of genes linked to these distal DA peaks revealed enrichment in trans-synaptic signaling and postsynaptic membrane regulation for LAMP5/LHX6 cells, consistent with fast and precise synaptic inhibition^37, 38, 39^ In contrast, LAMP5 cell–specific peaks were associated with pathways related to long-term synaptic depression, reflecting features of sustained inhibition in CGE-derived neurons (Extended Data Fig. 6b). Interestingly, the enrichment of ZIC4, USF1, and MEF2B motifs in primate LAMP5/LHX6 cells, but not in mice (Fig. 4b), suggesting an evolutionary divergence in the regulatory programs governing interneuron specialization. These transcription factors are known to regulate telencephalic regionalization, neuronal differentiation, and activity-dependent plasticity^40, 41, 42^. Their primate-specific motif enrichment therefore implies adaptive cis-regulatory innovations contributing to cortical expansion and refined inhibitory circuit modulation in primates. Moreover, the pronounced regional variation in the ratio of LAMP5/LHX6 to LAMP5 cells, with the highest abundance in the cingulate cortex (Fig. 4c), indicates potential region-specific functions. The preferential enrichment of LAMP5/LHX6 cells in association cortices may reflect their role in higher-order integrative and cognitive processing^43, 44^, whereas the broader distribution of LAMP5 cells across cortical areas likely supports more generalized inhibitory balance. By examining the shared CREs in LAMP5/LHX6 cells between macaques and humans, we found enrichment of CREs for genes in the pathways of monoatomic and inorganic cation transmembrane transport, such as *KCNQ3*, *SLC6A1* and *CACNA1G* (Extended Data Fig. 6c and 6d), suggesting the distinct role of this enriched cell type in the primate brain.

**Fig. 4.**
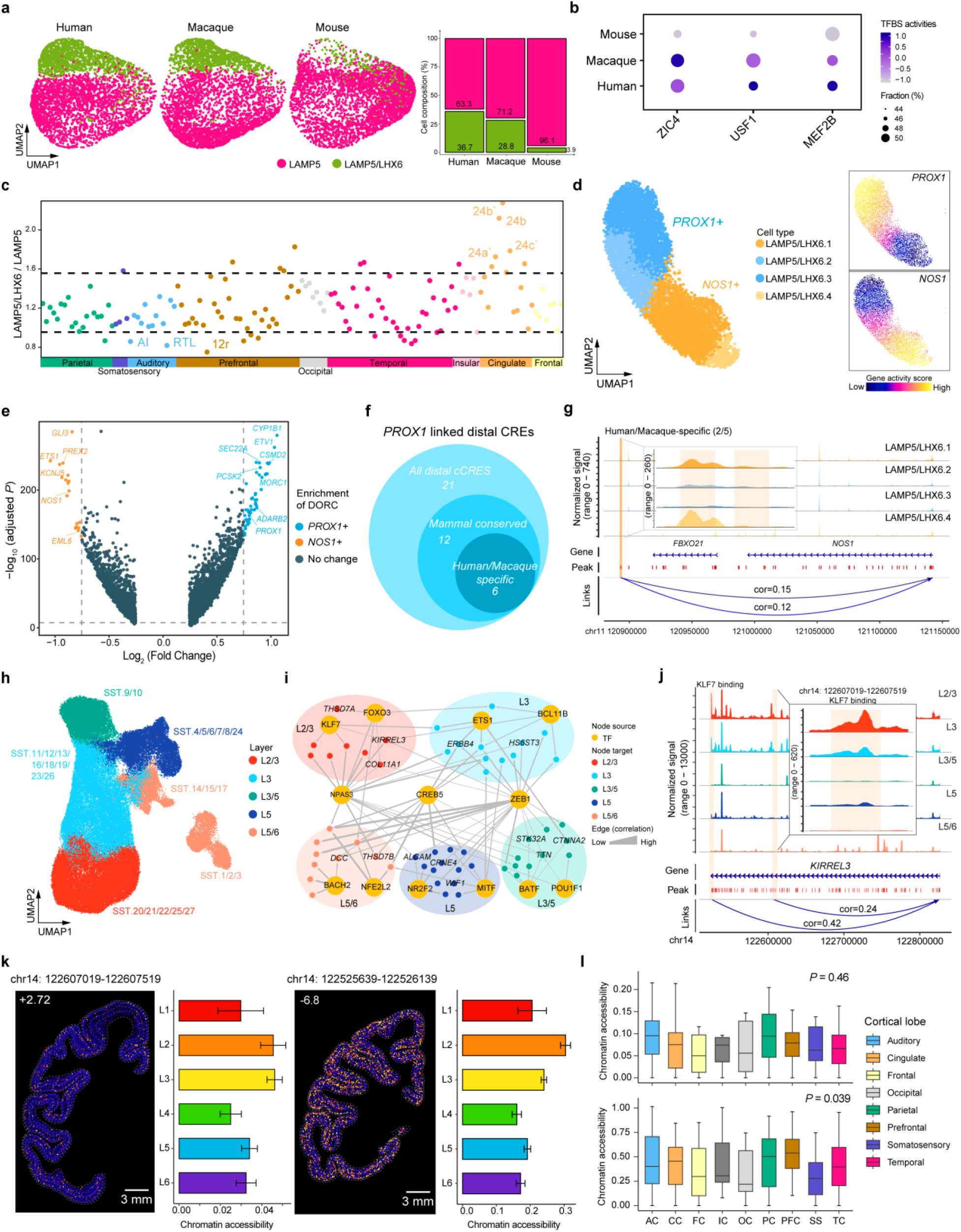
Characterization of specialized regulatory landscape of GABAergic neurons. **a**, The UMAP visualization (left) and relative percentage (right) of GABAergic LAMP5 and LAMP5/LHX6 cell subclasses among humans, macaques and mice. **b**, Dot plot showing the transcription factor binding site (TFBS) activities in LAMP5/LHX6 GABAergic neurons across species. **c**, Relative ratio of LAMP5/LHX6 vs. LAMP5 cell subclasses in various macaque cortical regions. **d**, UMAP embedding of LAMP5/LHX6 neurons (left) and normalized gene activity score of *PROX1* and *NOS1* (right). **e**, Volcano plot showing the differentially enriched DORCs in *PROX1*^+^ and *NOS1*^+^ populations of LAMP5/LHX6 neurons. Differential enrichment was defined as adjusted *P*-value < 1×10⁻⁵ and |log₂FC| > 0.75. **f**, The number of distal CREs linked to TSS region of *PROX1*. **g**, Genome browser track view showing two primate-specific distal CREs linked to TSS region of *NOS1* in LAMP5/LHX6 cell clusters. **h**, UMAP embedding of GABAergic SST neurons labeled with their layer specificity. **i**, Gene regulatory networks for layer-specific SST neuron groups. **j**, Genome browser track showing two differential accessible distal CREs linked to *KIRREL3* in L2/3 SST neurons. The two CREs harbor binding motifs of TF *KLF7*. **k**, Spatial visualization of chromatin accessibility of CREs linked to *KIRREL3* in SST neurons in representative sections at different coordinates in EBZ numbers (left). The bar plot illustrates the mean chromatin accessibility of CREs across different layers (right). Error bars represent standard error of the mean (SEM). **l**, Boxplot showing chromatin accessibility of two CREs linked to *KIRREL3* (upper/lower correspond to left/right in **k**) in SST neurons across cortical lobes. *P* values were calculated by Kruskal-Wallis test.

#### Further characterization of LAMP5/LHX6 GABAergic cell subpopulations

Examination of chromatin accessibility in LAMP5/LHX6 cells showed that these cells could be divided into two populations with differential accessibility of the *PROX1* gene (Fig. 4d), in line with our finding based on a previous transcriptomic dataset on primate cortical cells^21^ (Extended Data Fig. 6e). Moreover, we also found differential chromatin accessibility for the marker gene *NOS1* in macaque LAMP5/LHX6 cells (Fig. 4d), suggesting complementary roles of *PROX1* and *NOS1*. To validate the presence of *PROX1*⁺ and *NOS1*⁺ subpopulations within LAMP5/LHX6 cells, we analyzed a published MERFISH dataset from human dlPFC^45^, which confirmed the existence and spatial distribution of these two subgroups at single-cell resolution (Extended Data Fig. 6f-k). We then identified the domains of regulatory chromatin (DORC) in each of the LAMP5/LHX6 cell populations and found that DORC of *SEC22A*, *ADARB2* and *PROX1* genes were upregulated in *PROX1*^+^ cells, and DORC of *ETS1, NOS1* and *EML6* genes were upregulated in *NOS1^+^* cells (Fig. 4e and Extended Data Fig. 6l). This result further supports the importance of *PROX1* and *NOS1* in determining the heterogeneity of primate cortical LAMP5/LHX6 cells. Among the CREs associated with the *PROX1* and *NOS1* genes in LAMP5/LHX6 cells, 6 out of 21 CREs linked to *PROX1* were human/macaque-biased (Fig. 4f), whereas 2 out of 5 distal CREs linked to *NOS1* were human/macaque-biased (Fig. 4g). Differential spatial distributions were also observed, with the *PROX1^+^* population being enriched in layers 3, 5, and 6, while the *NOS1^+^* population was enriched in layers 2, 3, and 5 (Extended Data Fig. 6n). A previous study of human embryonic ganglionic eminence development^46^ identified an MGE-derived LHX6⁺CRABP1^-^ population proposed as progenitors of adult LAMP5/LHX6 neurons, which we found also co-express *PROX1* and *NOS1*. Consistently, in macaque, adult LAMP5/LHX6 neurons from our previously snRNA-seq dataset^21^ clustered with an embryonic *LHX6*⁺/*CCK*⁺ population (*CRABP1*^-^) reported as their putative progenitors^42^, likewise marked by *PROX1* and *NOS1* expression (Extended Data Fig.6n, o). These findings support a developmental continuity from MGE-derived embryonic progenitors to adult PROX1⁺ and NOS1⁺ LAMP5/LHX6 subpopulations.

#### Chromatin accessibility of CREs may underlie layer- and region-preference of SST neurons

SST cells were resolved into 27 transcriptionally distinct clusters characterized by differential chromatin accessibility (Extended Data Fig. 7a), as shown by the top differentially accessible (DA) peaks (Extended Data Fig. 7b). Genes linked to these DA peaks indicated cell type-specific epigenomic regulation. Notably, clusters enriched for high *HTR2A/HTR2B* gene activity (SST.1, SST.2 and SST.3) corresponded to serotonergic-responsive interneurons (Extended Data Fig. 7c). To further characterize functional differences, we performed rGREAT analysis on DA peaks from these three clusters and found that distinct SST subgroups were enriched in different biological processes: SST.1 in neuropeptide signaling pathway, SST.2 in energy homeostasis, and SST.3 in synapse organization (Extended Data Fig. 7d), underscoring the biological relevance of this fine-grained classification.

Previous study has revealed the layer-specific distribution of SST neurons that contributed to various anatomical and electrophysiological features of the human cortex^47^. Layer enrichment of various SST cell types was also found in single-cell spatial transcriptome map of the macaque cortex^21^. We then further focused on the layer-specific regulatory programs of various SST cell types. We classified layer-specific SST neurons based on clustering analysis of their chromatin accessibility profiles and annotated them based on their layer preferences (from L2/3 to L5/6, Fig. 4h), using gene activity scores of previously identified layer marker genes^21^ (Extended Data Fig. 7e). Further GRN analysis revealed that different layer-enriched SST cell types showed layer-enriched activity scores for genes targeted by different TFs, such as *KLF7* and *FOXO3* for L2/3 SST cells, *ETS1* and *BCL11B* for L3 SST cells, *BATF* and *POU1F1* for L3/5 SST cells, *NR2F2* and *MITF* for L5 cells, and *BACH2* and *NFE2L2* for L5/6 SST cells (Fig. 4i and Supplementary Table 5). As an example, we demonstrated that the preferential accessibility of two CREs associated with *KIRREL3* for KLF7 binding in L2/3 on the spatial transcriptome map (Fig. 4j, k). Interestingly, we also noticed that the accessibility of the two CREs showed regional differences, with the highest levels in parietal and prefrontal lobes, respectively (Fig. 4l). In another example, we found the preferential L5/6 accessibility of CREs associated with *DCC* for BACH2 binding (Extended Data Fig. 7f, g) and the highest level of this CRE accessibility in the prefrontal and cingulate cortical regions (Extended Data Fig. 7h). These findings support the notion that binding of specific TFs to accessible CREs for genes expressed in SST cells may determine their layer-specific and region-enriched distribution.

### Characterization of CREs for cortical non-neuronal cells

#### Cross-species comparison of various non-neuronal cell types

We have further explored the diversity of gene regulatory mechanisms that may be responsible for generating distinct non-neuronal cell types and their distribution in various macaque brain regions, as revealed by the previous spatial transcriptome analysis^21^. Cross-species comparison of our macaque data with those from humans and mice^11, 15^ showed that these non-neuronal cell types are all present in the cortices of the three species, with the percentage of various non-neuronal cells varied among the three species. While the astrocytes were found to be present at similar percentages, the percentages of oligodendrocytes (OLGs) increased gradually from mice to macaques and humans, and that of microglia in the human cortex was much higher than that found in both macaque and mouse cortices (Extended Data Fig. 8a, b). Notably, we found that the percentages of cortical oligodendrocyte precursor cells (OPCs) were similar between macaques and humans, but much higher than that in mice (Extended Data Fig. 8a, b). This finding is in line with the prolonged myelination and altered gene regulation in the oligodendrocyte lineage of the primate brain^2, 48, 49, 50^.

#### Cortical hierarchy-dependent regulation of OPCs and OLGs

We next examined the abundance of OPCs vs. OLGs across the 143 cortical regions and found the opposite abundance of OPCs and OLGs in both snATAC-seq and spatial transcriptome data (Fig. 5a and Extended Data Fig. 8c). Interestingly, we observed a strong negative correlation (*R* = -0.9, *P* < 2.2×10^-16^) between the percentages of OPCs and OLGs among various cortical regions in the visual system^51^ (Fig. 5b). Specifically, higher abundances of OPCs and OLGs were found in cortical regions of high and low hierarchy levels, respectively. The high potential for oligodendrocyte proliferation due to OPCs in regions of higher hierarchy suggests higher plasticity in the regulation of axon myelination in these regions. To further understand the differences in regulatory elements for OPCs and OLGs among different brain regions, we performed Spearman correlation analysis of chromatin accessibility and the hierarchy levels of various brain regions in the visual system. The results showed a consistent higher number of CREs exhibiting chromatin accessibility negatively correlated with the hierarchy level in OPCs regardless of thresholds for coefficients and *P* values (Extended Data Fig. 8d). Interestingly, more than 30% of these negatively correlated CREs were human/macaque-biased (Extended Data Fig. 8e, f), and their linked genes were significantly involved in GO terms including glutamatergic synapse, visual system development and positive regulation of kinase activity (Extended Data Fig. 8g and Supplementary Table 6). For examples, one mammal-conserved CRE showing open chromatin peaks only in OPCs was linked to *PDGFRA*, a typical marker gene for OPCs (Fig. 5c). Another example is a human/macaque-biased CRE with open chromatin only in OPCs that linked to *FBLN2*, which is an extracellular matrix inhibitor of OPC differentiation^52^. These results suggest the reciprocal regulation between OPCs and OLGs and the importance of OPCs in regulating higher cortical functions.

**Fig. 5.**
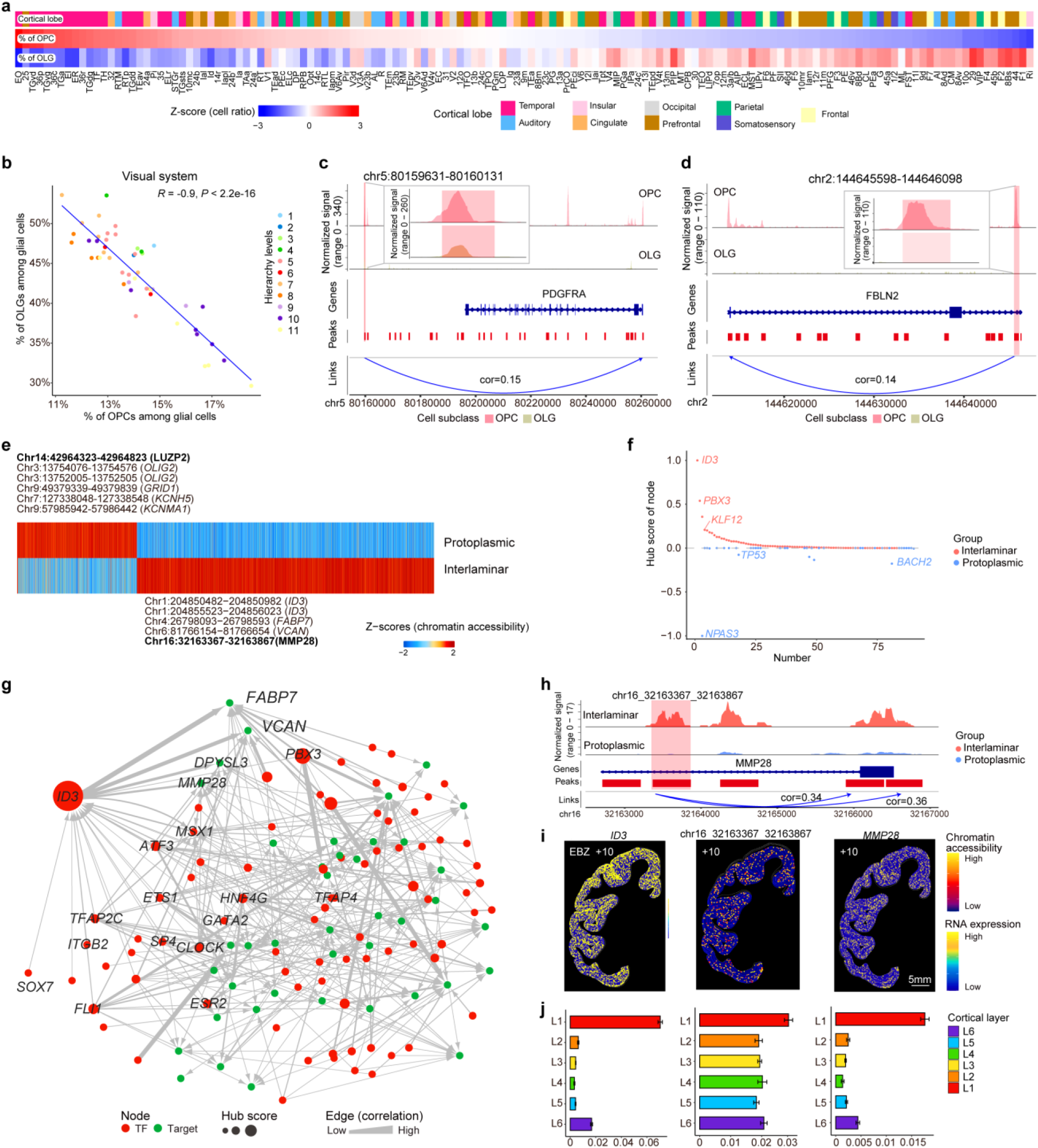
Regional specificity of CREs in cortical non-neuronal cells. **a**, Heatmap showing the percentages of OPCs and OLGs among all glial cells in various brain regions of the macaque cortex after spatial mapping. The brain regions are ordered by the decreasing percentage of OPCs. **b**, Correlation between the percentage of OPCs and OLGs in various cortical regions in the visual system. Brain regions are colored by their hierarchical levels. Correlation coefficients and *P* values were calculated by Spearman correlation analysis. **c**, Genome browser track view showing one mammal-conserved CRE linked to the TSS region of *PDGFRA* in OPCs. **d**, Genome browser track view showing one human/macaque-biased CRE linked to the TSS region of *FBLN2* in OPCs. **e**, Heatmap showing the chromatin accessibility of CREs in protoplasmic and interlaminar astrocytes. **f**, Dot plot showing the hub score of enriched TFs in the GRNs of interlaminar and protoplasmic astrocytes. The x-axis is ordered by the decreasing hub scores of TFs in the GRNs. **g**, Gene regulatory network of TFs and targeted genes in interlaminar astrocytes. **h**, Genome browser tracks showing the chromatin accessibility of CRE linked to *MMP28* between protoplasmic and interlaminar astrocytes. **i**, Spatial maps of expression patterns of *ID3* and *MMP28*, and chromatin accessibility of the CRE linked to *MMP28* at a representative section (EBZ +10). **j**, Bar plots showing quantitative levels of *ID3* and *MMP28* expression as well as chromatin accessibility of the CRE linked to *MMP28* across six cortical layers. Error bars represent standard error of the mean (SEM).

#### CREs-determined GRNs between interlaminar and protoplasmic astrocytes

Previous spatial transcriptome studies have shown laminar preferences of astrocytes (ASCs)^21, 45^. Our snATAC-seq-based cell clustering analysis further revealed differential chromatin accessibility of CREs associated with interlaminar ASCs (enriched in L1) and protoplasmic ASCs (enriched in L2-L6) (Extended Data Fig. 8h). For example, higher chromatin accessibility was found for CREs linked to *ID3*, *FABP7*, *VCAN* and *MMP28* and those linked to *LUZP2*, *GRID1*, *KCNH5* and *KCNMA1* in interlaminar and protoplasmic ASCs, respectively (Fig. 5e). Based on GO analysis, CREs found in the interlaminar ASCs were enriched in genes associated with “cell adhesion” and “cell projection” pathways, whereas those in the protoplasmic ASCs were associated with “intracellular sodium ion homeostasis”, “GABAergic synapse”, and “positive regulation of calcium ion transport” (Extended Data Fig. 8i). Moreover, we constructed GRNs for the two groups of ASCs and found that interlaminar ASCs showed high enrichment of hub TF genes *ID3* and *PBX3* (Fig. 5f). The targeted genes of *ID3* (including *GATA2*, *MSX1*, *TFAP2C* and *PBX3*) were also involved in the central nervous system development (Fig. 5g). On the other hand, the protoplasmic ASCs were enriched in the hub TF gene *NPAS3* (Fig. 5f), which contributes to neurodevelopmental transcription factor networks and the regulation of brain glucose metabolism^53^ (Extended Data Fig. 8j). As examples, the CRE linked to *MMP28* showed high chromatin accessibility in interlaminar but not protoplasmic ASCs (Fig. 5h). Spatial mapping of this CRE also showed higher chromatin accessibility in L1, in line with the high expression of its linked gene *MMP28* and its binding TF *ID3* in L1 (Fig. 5i, 5j and Extended Data Fig. 8k). In contrast, one of the NPAS3-targeted CRE that linked to *LUZP2* showed significantly higher chromatin accessibility in protoplasmic ASCs, and spatial mapping of this CRE also showed higher chromatin accessibility across cortical layers, consistent with spatial expression pattern of *LUZP2* (Extended Data Fig. 8l, m). Polymorphic variants in *LUZP2* have been reported to be associated with Alzheimer’s disease^54^, schizophrenia^55^, intelligence^56^, and verbal memory^57^. Taken together, these results provide new information on regulatory programs underlying the diversity and spatial organization of non-neuronal cells in the primate brain.

### Association of cell type-specific CREs with brain disease risk SNPs

#### LDSC analysis of cell type-specific CREs and brain diseases

To explore the contribution of cell type-specific *cis*-regulatory elements (CREs) to the genetic architecture of brain diseases, we applied linkage disequilibrium score regression (LDSC) to test the enrichment of GWAS-identified risk SNPs from 28 brain disorders and human traits in our cell type-specific CREs (Supplementary Table 7). Hierarchical clustering based on enrichment similarity revealed strong associations between neuronal CREs, particularly those in glutamatergic neurons, and multiple brain disorders (Fig. 6a, 6b and Extended Data Fig. 9a; Full list of cell types in Supplementary Table 7). For examples, schizophrenia (SCZ) and bipolar disorder (BP) risk SNPs were enriched in CREs of superficial-layer glutamatergic neurons, whereas BP SNPs were also present in PVALB and SST neurons. Generalized epilepsy (GE) SNPs were enriched in RELN neurons, and Alzheimer’s disease (AD) SNPs were concentrated in microglia (Fig. 6a). Furthermore, SNPs related to cognitive traits such as cognitive function (CGF), intelligence quotient (IQ), reaction time (RT), and years of education (YED) showed the highest enrichment in L4 glutamatergic neurons (Fig. 6a). These results highlight the selective localization of disease-associated variants in CREs of specific neuronal subtypes, underscoring their potential role in disease susceptibility.

**Fig. 6.**
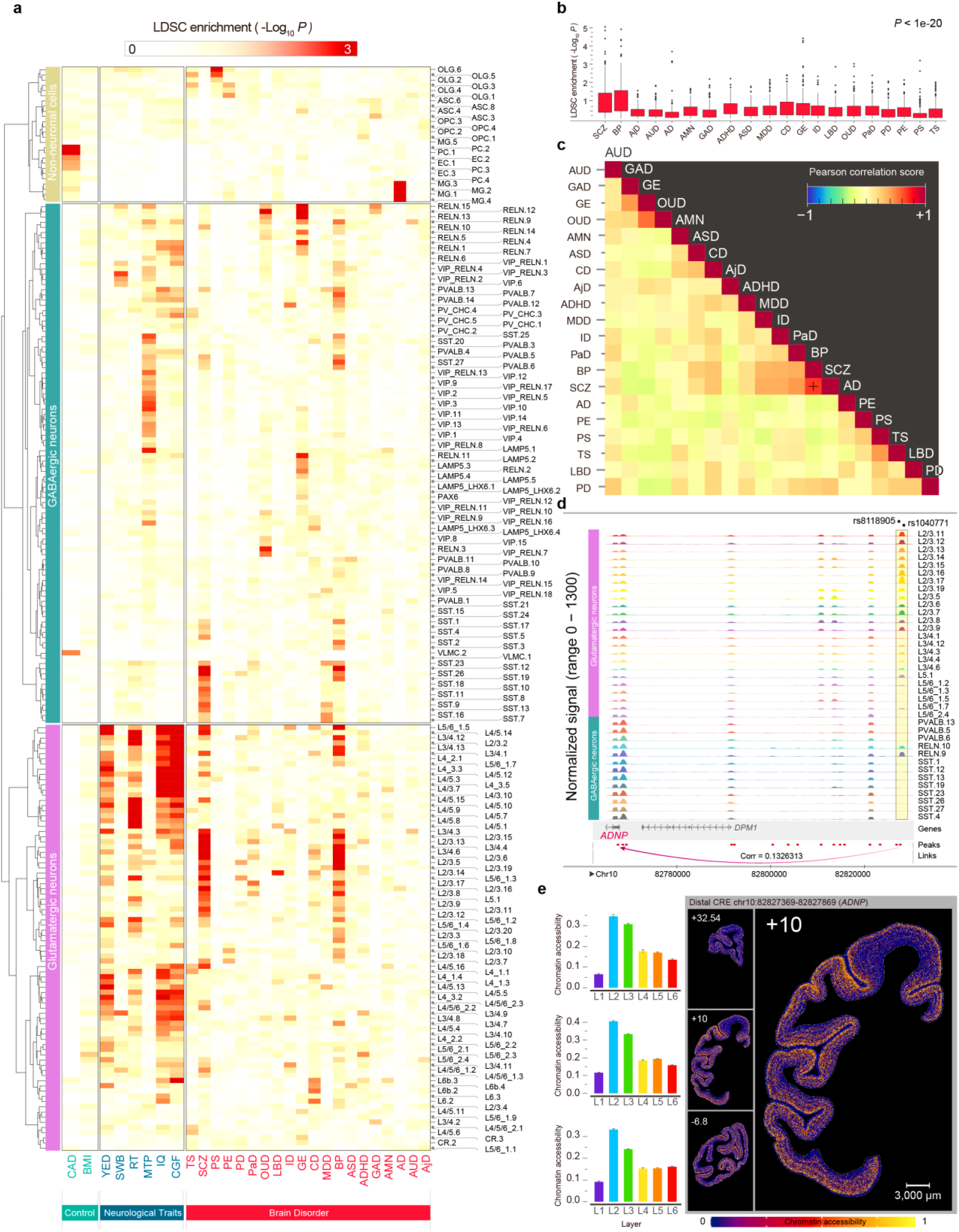
Association of cell type-specific CREs with risk SNPs of various brain disorders and human traits. **a**, Heatmap showing the enrichment of cell type-specific CREs containing risk SNPs of various brain disorders and human traits for all neuronal and non-neuronal cell types from snATAC-seq data of the macaque cortex. Intensity of the red color codes LDSC enrichment score (defined as -log10 *P* values, with *P* representing the probability of CREs containing risk SNPs). Black triangles mark the two brain disorders with most prominent enrichment probabilities. AjD: adjustment disorder, AUD: alcohol dependence, AD: Alzheimer disease, AMN: amnesia, GAD: anxiety disorder, ADHD: Attention deficit hyperactivity disorder, ASD: Autism spectrum disorder, BP: bipolar disorder, MDD: Major depression disorder, CD: cognitive disorder, GE: generalized epilepsy, ID: Insomnia, LBD: Lewy body dementia, OUD: opioid dependence, PaD: panic disorder, PD: Parkinson Disease, PE: partial epilepsy, PS: psychosis, SCZ: Schizophrenia, TS: Tourette syndrome, CGF, IQ: Intelligence, cognitive function, MTP: Morning Person, RT: reaction time, SWB: subjective well-being, YED: Years of Education, BMI: Body mass index, CAD: coronary artery disease. **b**, Boxplot showing the relative enrichment levels across different diseases. Significant difference was observed among different diseases (*P* < 1e-20, Kruskal-Wallis test). **c**, Heatmap showing the Pearson correlation coefficients for the association of cell type-specific CREs with disease risk SNPs loci for various brain disorders. Star marked the high similarity in disease risk SNP-associated cell type-specific CREs between schizophrenia and bipolar disorder. **d**, Genome browser tracks showing chromatin accessibility of CREs for gene *ADNP* that contain risk SNPs (marked by dots) of both SCZ (schizophrenia) and BP (bipolar disorder). The arrow line indicates the targeted gene *ADNP*. **e**, Spatial maps of cell type-enriched CREs containing SCZ- and BP-associated SNPs loci shown in **d** at three representative sections along the coronal coordinates, indicating the layer-specific distribution of the CRE for *ADNP*.

#### CREs associated wi th SCZ and BP risk SNPs

We next examined the genomic context and evolutionary properties of CREs colocalized with disease-associated SNPs. Approximately 40% of these CREs were intronic, and ∼50% were located more than 1 kb from transcription start sites (Extended Data Fig. 9b). Compared to background peaks, they exhibited significantly lower phyloP scores^58, 59^ (Extended Data Fig. 9c), suggesting rapid evolutionary turnover. Enrichment patterns across diseases showed the highest correlation between SCZ and BP (Fig. 6c and Extended Data Fig. 9d), consistent with their known genetic overlap^60^. Notably, we identified two shared SNPs (rs8118905 and rs1040711) within a CRE linked to the *ADNP* promoter^61, 62^, with accessibility peaking in L2/3 glutamatergic neurons (Fig. 6d). Spatial mapping of this CRE also showed the highest level of accessibility in L2 and L3 of the cortex (Fig. 6d and 6e). In addition, we found two other SNPs rs2051307 and rs7753616 were linked to *SETBP1* gene in BP and *MDGA1* gene in SCZ, respectively (Extended Data Fig. 9e). Both genes were related to neurodevelopment^63, 64^, and the spatial distribution of SNP-associated CREs for the two genes were also enriched in the L2/3 glutamatergic neurons (Extended Data Fig. 9f). Together, these findings indicate that SCZ- and BP-associated SNPs preferentially disrupt regulatory elements active in L2/3 glutamatergic neurons, highlighting a shared neurodevelopmental basis for disease risk.

#### Human/macaque-biased CREs harboring disease risk SNPs

To assess the evolutionary specificity of disease-associated CREs, we classified SNP-containing CREs as human/macaque-biased or mammal-conserved. Glutamatergic neurons, especially the L4_3 subclass, harbored significantly more human/macaque-biased disease-associated CREs than other cell types (Fig. 7a). In L4_3 subclass, human/macaque-biased CREs carrying SCZ SNPs displayed higher accessibility than mammal-conserved CREs (Fig. 7b). Among them, genes linked to human/macaque-biased CREs (181 gene; including *ABCA7^65^*, *ADORA1^66^* and *CHGA^67^*) and mammal-conserved CREs (293 genes, including *CHRM4^68^*, *CXXC1^69^* and *OPCML^70^*) showed only minimal overlap (12 genes, including *PDYN)^71^*(Fig. 7c and Extended Data Fig. 9g). Spatial mapping showed cell type- and region-specific accessibility patterns of these CREs, with L4_3.11 neurons in the occipital lobe exhibiting the highest accessibility (Fig. 7d). Thus, SCZ-associated regulatory mechanisms appeared to be highly cell type- and brain region-specific. Gene regulatory network analysis of L4_3 neurons identified transcription factors CTCF*^72^,* NEUROD6*^73^,* SREBF1^74^, HIF1A^75^ and CXXC1^69^ as central regulators (Fig. 7e). Furthermore, the human/macaque-biased CRE linked to *GFRA2^76^* and the mammal-conserved CRE linked to *NEFH^77^*, containing binding sites for regulators of SREBF1 and HIF1A, respectively, both exhibited the highest chromatin accessibility in L4 of the spatial maps (Fig. 7f and Extended Data Fig. 9h). Collectively, these results indicate that primate-evolved regulatory networks may play a pivotal role in SCZ pathology.

**Fig. 7.**
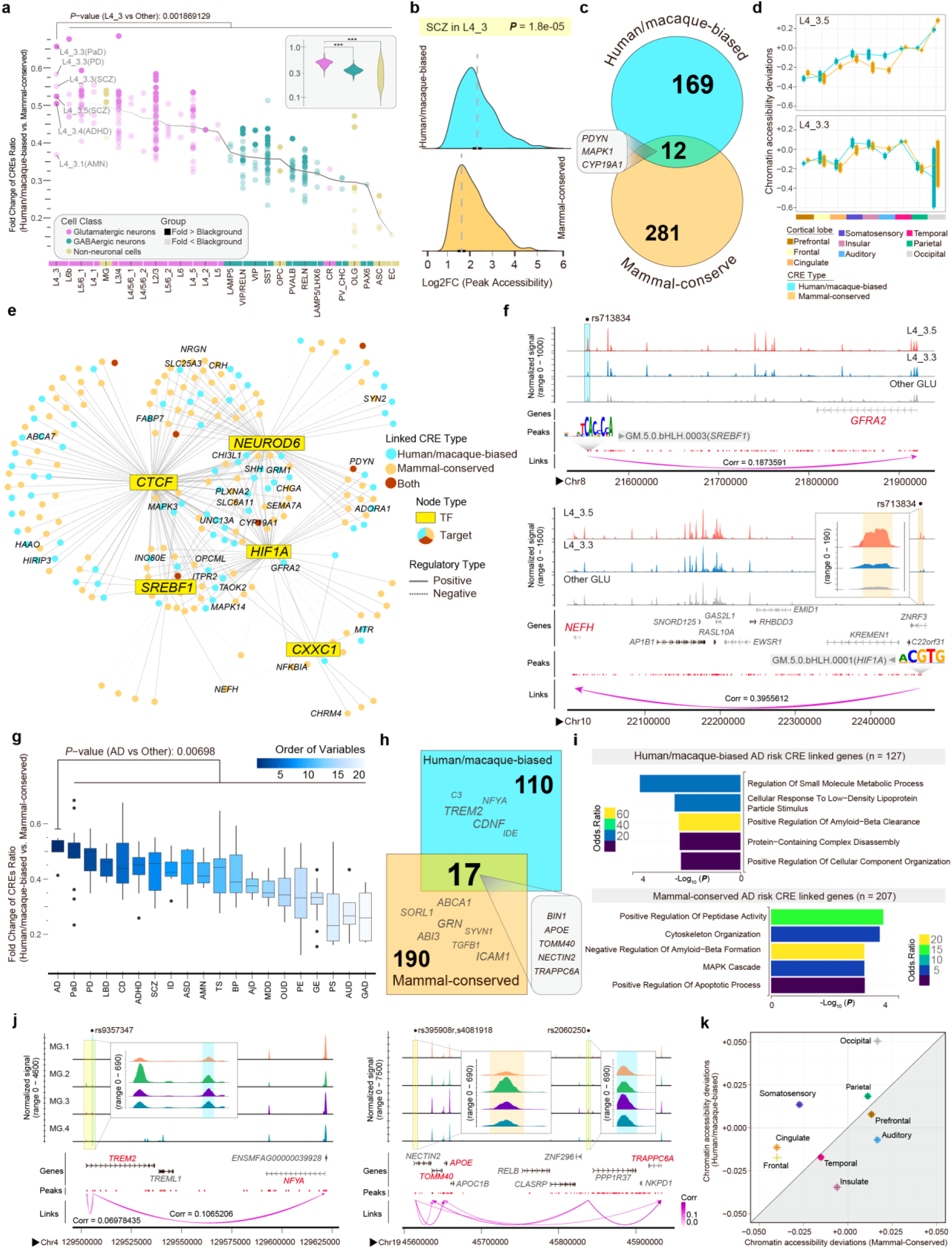
Prominent association of human/macaque-biased CREs with risk SNPs of various brain disorders. **a**, Dot plot showing the ranked distribution of human/macaque-biased vs. mammal-conserved CRE ratios (with disease-risk SNPs) across cell subclasses. Cell types with fold changes below the background are shown with reduced transparency. Fold changes for glutamatergic, GABAergic, and non-neuronal cell types are showed in the top-right corner. Significant difference was observed between L4_3 glutamatergic neuron and other cell subclasses (*P* = 0.001869, Wilcoxon rank-sum test). The violin plot (upper left) illustrates comparisons between glutamatergic neurons, GABAergic neurons, and non-neuronal cells, with asterisks indicating nominally significant differences (Wilcoxon rank-sum test, *** *P* < 0.001). **b**, Ridge plot depicting the chromatin accessibility of human/macaque-biased and mammal-conserved CREs (colocalized with SCZ risk SNPs) in the L4_3 glutamatergic cluster. Significant difference was observed between these two groups (*P* = 1.8e-05, T-test). **c**, Venn Diagram showing the overlap of genes associated with human/macaque-biased and mammal-conserved CREs (colocalized with SCZ risk SNPs) in the L4_3 glutamatergic cluster. **d**, Bar plot (Mean ± SE) illustrating the relative chromVAR deviations of SCZ risk SNPs associated CREs across cortical lobes in the L4_3 glutamatergic cluster. **e**, Gene regulatory network (GRN) showing the key transcription factors (TFs) for targeted genes linked to CREs associated with SCZ risk SNPs in L4_3 neuron. SCZ ontology genes were labeled. **f**, Genome browser tracks showing chromatin accessibility in L4_3 neurons at SCZ risk SNP-associated CREs linking to *GFRA2* and *NEFH*, respectively. Highlighted regions are marked with color bands. **g**, Box plot showing the ranked distribution of the fold change in the relative composition of human/macaque-biased versus mammal-conserved CREs (colocalized with disease risk SNPs) across various brain disorders. Significant difference was observed between Alzheimer’s disease (AD) and other brain disorders (*P* = 0.00698, Wilcoxon rank-sum test). **h**, Venn Diagram illustrating the overlap of genes associated with human/macaque-biased and mammal-conserved CREs (colocalized with AD risk SNPs) in microglia cells. **i**, Pathways enriched by genes linking to AD risk SNP associated CREs in microglia. **j**, Genome browser tracks plot showing chromatin accessibility in microglia at the distal regions of genes linking to the AD risk SNP associated CREs: *TREM2* and *NFYA* (left, primate-specific), as well as *APOE*, *TOMM40*, and *TRAPPC6A* (right, human/macaque-biased and mammal-conserved). Highlighted regions are marked with color bands. **k**, Dot plot illustrating the relative chromVAR deviations of human/macaque-biased and mammal-conserved CREs (colocalized with AD risk SNPs) across brain lobes in microglia.

We also investigated the distribution and functional properties of human/macaque-biased and mammal-conserved CREs carrying AD risk SNPs in microglia. AD SNP–associated CREs were exclusively localized to microglia (Fig. 6a), with the highest ratio of human/macaque-biased to conserved CREs observed among all brain diseases examined (Fig. 7g). These two CRE classes were largely distinct in their target genes (127 vs. 207 genes with 17 shared) (Fig. 7h). Human/macaque-biased CREs carrying AD risk SNPs were linked to genes involved in pathways such as amyloid-beta clearance, LDL response, and small molecule metabolism—absent from those associated with conserved CREs (Fig. 7i). Notable AD-related genes linked to human/macaque-biased CREs included *C3^78^*, *TREM2*^79^, *NFYA^80^*, *CDNF^81^* and *IDE^82^*. For instance, one human/macaque-biased CRE in microglia was linked to *TREM2* and *NFYA.* On the other hand, both mammal-conserved and human/macaque-biased CREs were linked to AD-related genes^83, 84, 85^ *APOE, TOMM40* and *TRAPPC6A* in microglia (Fig. 7j). Spatial accessibility analysis revealed that human/macaque-biased CREs were most accessible in occipital and somatosensory cortices, whereas conserved CREs peaked in insular and auditory cortices (Fig. 7k). These results suggest that human/macaque-biased microglial CREs regulate distinct gene programs relevant to AD pathogenesis, highlighting their potential as species-specific therapeutic targets.

#### Differential enrichment of disease SNPs in human- and macaque-specific CREs of LAMP5/LHX6 GABA neurons

Given that LAMP5/LHX6 neurons are enriched in both primates (Fig. 4a), we next assessed whether disease risk SNPs are differentially enriched between human-specific and macaque-specific CREs in this cell type. We focused on three brain disorders—attention deficit hyperactivity disorder (ADHD), cognitive disorder (CD), and generalized epilepsy (GE)—which exhibited significant heritability enrichment in our LDSC analysis (Fig. 6a). For each trait, genes linked to CREs harboring risk SNPs showed limited overlap between the two species (human vs. macaque, ADHD: 919 vs. 1246 genes, 65 shared; CD: 771 vs. 1055 genes, 70 shared; GE: 470 vs. 634 genes, 33 shared). Notably, within conserved genomic regions, disease-associated SNPs mapped to distinct CREs that were either human- or macaque-specific (Extended Data Fig. 10a-c). For examples, an ADHD risk SNP (rs34376065) was co-localized with a human-biased CRE linked to the *NTNG1* gene. In contrast, open chromatin peaks were only found in the macaque genome for CREs harboring several ADHD risk SNPs (Extended Data Fig. 10b). These results highlight species-specific divergence in regulatory architectures underlying disease heritability, even within consensus primate cell types such as LAMP5/LHX6 neurons.

## Discussion

We have performed a comprehensive study of chromatin accessibility at the single-cell level across 142 brain regions of the macaque cortex and identified specific CREs for various cell types. Leveraging the newly developed method for integrating chromatin accessibility and spatial transcriptome data, we found that many cell type-specific CREs showed prominent laminar and regional preferences in their chromatin accessibility, thus providing gene regulatory mechanisms underlying the laminar and regional diversification of cell types. Cross-species comparison with human and mouse snATAC-seq data further identified CREs associated with brain disease-related genes in human/macaque-enriched L4 glutamatergic and GABAergic cell types. Further comprehensive association analysis between cell type-specific CREs and brain disease risk SNPs revealed potential contribution of human/macaque-biased CREs in various brain diseases. Notably, human/macaque-biased CREs in L4 glutamatergic neurons are much more prominently associated with risk SNPs for schizophrenia. Furthermore, we also identified human/macaque-biased CREs that regulate AD pathogenesis-related genes (*C3*, *TREM2*, *NFYA*, *CDNF* and *IDE)* exclusively in microglia. This study offers a comprehensive database for further investigating gene regulatory mechanisms underlying cell type diversity in the primate brain as well as brain disease pathogenesis.

Our analysis of the chromatin accessibility landscape in primate and mouse cortices revealed the role of human/macaque-biased CREs in the emergence of human/macaque-biased cell types as well as their potential involvement in psychiatric diseases. For example, in human/macaque-biased L4 glutamatergic neurons, we identified CREs localized in the regulatory region of *DCC*, a gene involved in neurodevelopment and psychiatric disorders^86, 87, 88^. We also found that human trait-related SNPs largely resided in CREs of human/macaque-biased glutamatergic cell types, suggesting potential contributions of these neuronal types to cortical development underlying human intellectual traits. The discovery of distinct presence of human/macaque-biased CREs related to some brain disorders such as schizophrenia and AD supports the notion that these brain diseases are due to cortical pathology that are largely primate-specific.

Previous studies have shown that cortical LHX6^+^LAMP5^+^ GABAergic neurons are more abundant in primates^4^, but their molecular diversity and regulatory features remain poorly characterized. Here, we further subdivided the LAMP5^+^LHX6^+^ population into two transcriptionally and epigenetically distinct subtypes—*PROX1*⁺ and *NOS1*⁺—characterized by mutually exclusive chromatin accessibility at their respective lineage-defining genes (*PROX1* for CGE origin^46^, *NOS1* for MGE origin^89^), as well as distinct laminar distributions within the cortex. Beyond previous transcriptomic observations, our analysis revealed subtype-specific DORCs and identified multiple human/macaque-biased CREs associated with *PROX1* and *NOS1*, suggesting evolutionary innovation in the regulatory architecture of these interneurons. These findings highlight previously unrecognized epigenetic and spatial heterogeneity within LAMP5⁺LHX6⁺ neurons and provide a resource for understanding their developmental origin and primate-specific specialization.

Our comprehensive analysis of chromatin accessibility and spatial transcriptome allowed us to identify differences in chromatin accessibility across various cortical regions. This information is critical for understanding the gene regulation underlying regional differentiation of cell types. Our finding of the reciprocal relationship between the abundance of OPCs and OLGs across all brain regions suggest that OLGs may provide negative regulation of OPC proliferation. The opposite trend in the abundance of OPCs and OLGs at different hierarchy levels of the visual system^51^ further suggests the functional role of axon myelination in neural properties at various hierarchy levels, and OPC biogenesis and myelination is tightly regulated across cortical hierarchy. The higher abundance of OPCs in cortices of higher hierarchy suggests their higher capability of generating new OLGs, contributing to the plasticity of myelin formation and regeneration in these cortices.

Elucidation of regulatory mechanisms underlying diversity of cortical cell types and their distribution is important for understanding not only the structure and function of the brain but also pathogenic mechanisms underlying brain disorders. Through association analysis between cell-type specific CREs and brain disease risk SNPs, we uncovered hundreds of disease-related CREs in the primate brain, including those regulating genes involved in SCZ, BP and AD. Our finding on AD risk SNP-associated CREs exclusively in microglia provides strong support to the notion that microglia play a critical role in the AD etiology^90, 91^. Notably, prominent AD risk SNP alleles were found in CREs linked to *TREM2* and *APOE* genes in microglia, and CREs for *TREM2* were found to be human/macaque-biased. In addition, SCZ- and BP-associated CREs were generally enriched only in the primate L2/3 glutamatergic neurons. Importantly, we found a large increase in the frequency of transposon element insertion in human/macaque-biased CREs. This suggests that the high frequency of brain disorder risk SNPs that are linked to human/macaque-biased CREs might be attributed to the extensive proliferation of cortical neurons during primate brain development.

## Methods

### Ethics statement

Animal protocol was reviewed and approved by the Biomedical Research Ethics Committee of CAS Center for Excellence in Brain Science and Intelligence Technology, Chinese Academy of Sciences (ION-2019011). Animal care complied with the guideline of this committee. Adult macaque monkeys (M. fascicularis; #1, 6-year-old, 4.2 kg, male; #2, 4-year-old, 3.7 kg, male; #3, 12-year-old, 5.1kg, female) were used in this study.

### Tissue collection and nucleus extraction

We sampled 128 cortical regions from Macaque #1 and 132 regions from Macaque #2 for snATAC-seq data generation. For Macaque #1, samples were collected from thick adjacent sections to the ones used for Stereo-seq. For Macaque #2, we dissected brain samples from block-face cortical regions of the hemisphere opposite to that used for Stereo-seq. Using tissue punchers (2.5 - 4 mm in diameter), we segmented the cortical areas at a depth of 1 – 2 mm in the cryostat. After dissection, the samples were promptly frozen in liquid nitrogen and then stored in dry ice or a -80 ℃ refrigerator. Throughout the sampling process, great care was taken to transfer the tissues to pre-chilled tubes without allowing them to thaw. The single nucleus suspension was prepared following the method described previously^92^. The isolated single nuclei were evenly split for snRNA-seq and snATAC-seq library preparation, respectively. In brief, frozen sections of monkey brain tissue were placed in a Dounce homogenizer containing 2 ml of pre-chilled homogenization buffer (containing 20mM Tris pH8.0, 500mM sucrose, 0.1% NP-40, 0.2U/μL RNase inhibitor, 1x protease inhibitor cocktail, 1% bovine serum albumin (BSA), and 0.1mM DTT), and the Dounce homogenizer was kept on ice during the grinding process. The tissue was homogenized with 10-15 strokes of pestle A, followed by 10-15 strokes of pestle B. Subsequently, 2 ml of homogenization buffer was added to the Dounce homogenizer, and the homogenate was filtered through 30 μm MACS SmartStrainers (Miltenyi Biotech, #130-110-915) into a 15 ml conical tube. The tube was then centrifuged at 500 g for 5 minutes at 4℃ to pellet the nuclei. The pellet was resuspended in 1.5 ml of blocking buffer (containing 1% BSA and 0.2U/μL RNase inhibitor in 1x PBS) and centrifuged again at 500 g for 5 minutes at 4℃ to pellet the nuclei. The nuclei were finally resuspended in cell resuspension buffer (PBS containing 1% BSA) for subsequent snATAC-seq library preparation.

### Library preparation for single-nucleus ATAC-seq

For the preparation of single-nucleus ATAC-seq libraries, we employed the DNBelab C Series Single-Cell ATAC Library Prep Set (BGI-research) with the procedure of step transposed, droplet encapsulation, pre-amplification, emulsion breakage, then the captured beads were collected. After DNA amplification and purification, the indexed sequencing snATAC library was already to sequence. Finally, we used the Qubit ssDNA Assay Kit (Thermo Fisher Scientific) to measure the concentrations of the sequencing libraries. All libraries were sequenced using a paired-end sequencing scheme by the ultra-high-throughput DIPSEQ T1 sequencer platform using the following read lengths: 70 bp for read 1, 50 bp for read 2, and 20 bp for the sample index sequencing scheme of the China National GeneBank (CNGB).

### Data processing and quality control of snATAC-seq data

We used the newest pipeline to process the snATAC-seq data of DNBelab C4 (https://github.com/MGI-tech-bioinformatics). First, the raw reads were separated into insertions and barcodes and then filtered and demultiplexed using PISA (v1.1) with a minimum sequencing quality of 20. Next, the filtered reads were aligned to the *Macaca fascicularis* genome (5.0.91) using Chromap (v0.2.3_r407) and mismatched barcode was not allowed. Reads with mapping quality less than 10 and reads mapped to the mitochondria or genome scaffold (chrAQ*, chrU*, chr*_random*, and chrK*) were filtered out and PCR duplicates were removed according to the cell barcode and mapping loci. The resulting BAM files were processed using d2c (v1.4.4) to identify barcodes from the same cell. The retained fragments of each library were used for further analysis.

Basic processing of the snATAC-seq data was performed using ArchR^93^ (v1.0.2) package in R (v4.2.2). Genome annotation file of *Macaca fascicularis* was download from Ensembl (release 91). Fragments files were loaded by “createArrowFiles” function, with chrM and chrY chromosomes excluded. Each sample was separately computed for counts for each tile per cell via “addTileMatrix” function with tileSize = 500. Doublets were identified by “addDoubletScores” function with Doublet score < 0.15. A total number of 1,615,430 cells from 405 snATAC-seq libraries was kept with threshold of TSS enrichment >= 5 and <=30 and Promoter ratio >= 0.05 and <= 0.6.

### Dimension reduction and clustering of snATAC-seq data

First, we used “addGroupCoverages” function and “addReproduciblePeakSet” function in MACS2^24^ (v2.2.7.1) to create a reproducible merged peak set for each library. A new matrix was created within the new merged peak set with “addPeakMatrix” function for downstream analyses. Next, we further performed an iterative LSI dimensionality reduction via the “addIterativeLSI” function with varFeatures = 50000. Harmony was applied to remove the batch effects across libraries. In order to better cluster the data, we applied a three-step clustering approach to refine cell type identification based on chromatin accessibility. This approach resulted in 3 classes, 29 subclasses and 230 cell types overall. We first carried out unsupervised clustering for all cells with resolution = 3, categorized the initial cell clusters into 3 categories based on classical marker genes, including glutamatergic neurons (SLC17A6, SLC17A7), GABAergic neurons (*GAD1*, *GAD2*) and non-neuronal cells (*GFAP*, *PDGFRA*, *PLP1*, *PLAC8*, *CLDN5* and *COL1A1*), then repeated the above analysis workflow with higher resolution for each category.

A total of 230 cell clusters were kept after removing clusters with the number of cells less than 100, or ambiguous clusters (expressing both glutamatergic (*SLC17A6*, *SLC17A7*) and GABAergic (*GAD1*, *GAD2*) marker genes or non-neuronal marker genes). We have provided a comprehensive list of marker genes used for the identification of ambiguous clusters in Supplementary Table 2. Previous study showed that most doublets would either form separate clusters, or would tend to end up at the fringes of other clusters (in graph embeddings)^94^. Therefore, we also applied the same procedure to further remove doublets from the clustering results. Specifically, cells were marked as doublets with their distances to the cluster center lower or higher than 1.5*interquartile range (25th and 75th percentile).

### Correspondence of cell types defined by snATAC-seq and snRNA-seq data

We calculated gene activity score for each gene by summing up the counts within the gene body + 2 kb upstream of our snATAC-seq data using ArchR “addGeneScoreMatrix” function, and compared with the snRNA-seq data derived from a recently published macaque cortex study^21^. We randomly selected 200 cells from each cell cluster for both snATAC-seq and snRNA-seq data, and then performed data normalization using “NormalizeData” function. Integration analysis was conducted with Seurat “IntegrateData” function with top 3000 repeatedly variable genes as features. The proportion of nuclei overlap between omics represents the number of cells clustered in the same clusters of the integrative clustering result divided by the total number of cells for specific cell types from the two omics datasets, using the "compare_cl" function of the published pipeline^6^.

### Identification of reproducible peak sets

We performed peak calling according to the ENCODE ATAC-seq pipeline (https://www.encodeproject.org/atac-seq/). Across each cell type, we pooled all appropriately paired reads to produce a pseudo-bulk ATAC-seq dataset for each biological replicate using “getFrags” function. In addition, we created two pseudo-replicates by dividing the reads from each biological replicate in half. We called peaks for each of the four datasets and a pool of both replicates independently. Peak calling was performed on the Tn5- corrected single-base insertions using MACS2 (v2.2.7.1) with these parameters: ––shift -75 --extsize 150 --nomodel --call-summits --SPMR -q 0.01. Finally, we extended peak summits by 250 bp on each side, resulting in a final width of 501 bp, for merging and downstream analysis. To identify a list of reproducible peaks, we filtered peaks with the following two criteria as in previous studies^15^, (1) detected in the pooled dataset and overlapped ≥50% of peak length with a peak in both individual replicates; or (2) detected in the pooled dataset and overlapped ≥50% of peak length with a peak in both pseudo-replicates.

Due to the nature of the Poisson distribution test in MACS2, the MACS2 score was influenced by the number of nuclei or the read depth. Thus, we utilized the adjusted normalization method which converted the MACS2 scores (−log_10_ (q-value)) to “score per million (SPM)”, and filtered reproducible peaks by choosing a SPM cut-off of 2. Finally, we only kept reproducible peaks on chromosome 1–19 and both sex chromosomes, and filtered blacklist regions. A union peak list for the whole dataset was obtained by merging peak sets from all cell types using Bedtools (v2.29.1).

### Chromatin co-accessibility identification

We used R package Cicero^26^ (v1.20.0) to compute co-accessible regions for all open regions. In each cell type, we randomly sampled 5,000 nuclei (All cells were retained for cell types with less than 5,000 nuclei). We set parameters of Cicero as: k = 50, window size = 500 kb, distance constraint = 250 kb. To avoid using an empirical cut-off, we also generated a random shuffled CRE-by-cell matrix and computed co-accessible regions on this shuffled matrix. Then, we fitted the distribution of co-accessibility scores from randomly shuffled background into a normal distribution model by using the R package fitdistrplus. Next, we tested each co-accessibility pair and set the cut-off at co-accessibility score with significance threshold of FDR < 0.01 to filter out co-accessible pairs.

We defined the CREs outside of ±1 kb of the gene transcriptional start site (TSS) as distal and the others as proximal. Then, we divided all co-accessible pairs into three groups: proximal-to-proximal, distal-to-distal, and distal-to-proximal. We focused only on distal-to-proximal pairs because they represent potential gene expression regulatory relationships. We classified all distal CREs in the above distal-to-proximal co-accessible pairs as final CREs.

We annotated the genome distribution of a total of 615873 CREs by “annotatePeak” function from R package ChIPseeker^95^ (v.1.34.0), setting 1kb upstream and 1kb downstream of the TSS as promoter.

### Differentially accessible CREs among different cell types and transcription factor (TF) binding motif analysis

We applied Wilcoxon rank sum test using the ‘‘getMarkerFeatures’’ function for identification of marker peaks with useMatrix = "PeakMatrix" at the cluster level. The CREs with log2 fold-change > 1 and FDR < 0.05 were identified as significant marker features for each cell type. Then, we removed the duplicated CREs and calculated the differential motifs across cell types with gimme maelstrom^96^. The “gimme.vertebrate.v5.0” was used as reference database.

We assessed TF binding motif enrichment in accessible peaks using chromVAR^97^ (version 1.24.0). Briefly, we initially employed the “addMotifAnnotations” function in ArchR to determine motif presence in the peak set. Subsequently, GC bias was corrected using “addBgdPeaks”, followed by calculation of deviation z-scores for each TF motif in each cell using the “addDeviationsMatrix” function. The cell-by-motif matrix was then stored in ArchR arrow files.

### Gene regulatory network inference

We used Python package CellOracle^28^ to infer GRNs by integrating with our previously published snRNA-seq data^28^. The snRNA-seq data were processed by Seurat with default parameters, using the ‘FindVariableFeatures’ to select the top 3,000 highly variable genes in each corresponding cell subclass, and randomly sampled 5,000 RNA nuclei for each cell subclass. After that, we predicted GRNs by three steps. First, we defined the co-accessible distal-to-proximal pairs as described above. Second, we mapped distal CREs to known motifs by the “scan” function of GimmeMotifs, using the macaque genome macFas5 and the “gimme.vertebrate.v5.0” motif database as references. Finally, we identified the regulatory relationships between TFs and the potential target genes by fitting a regularized linear regression model using processed snRNA-seq data. We evaluated key TFs potentially involved in regulatory functions across different cell types based on “eigenvector_centrality” values. Subsequently, we filtered downstream target genes highly regulated according to their “coef_abs” values. To identify hub TFs in the putative gene regulatory networks, we then calculated the hub score to infer the importance of different TFs. The hub score was calculated by “hub_score” function from R package igraph (v 2.1.1), with "coef_abs" value as a weight edge attribute

### Integration of snATAC-seq and Stereo-seq data

#### Data preprocessing

To enable the integration of the query modality (Stereo-seq) and the reference modality (snATAC-seq), a systematic data preprocessing pipeline was designed. First, gene names from both modalities were extracted, and their intersection was computed to generate a shared gene set. The data were subsequently trimmed to a unified gene feature space. For snATAC-seq data, the gene score matrix was subjected to normalization and logarithmic transformation for each cell. Subsequently, the top 3,000 highly variable genes were selected to construct a high-information gene subset, which reduced data dimensionality while retaining critical biological features.

To simulate the complexities and uncertainties inherent in cross-modal data integration and to create a more challenging and generalizable data distribution, the gene score matrix of snATAC-seq was intentionally perturbed with added noise. During the perturbation process, a disturbance matrix p_ik_ was constructed based on the proportional calculation and normalization of gene expression values. This matrix comprised two components: gene-level perturbation and cell-level perturbation.

Gene-level perturbation was calculated based on the ratio of average gene expression values between the Stereo-seq and snATAC-seq modalities. Let X_ik_ denote the gene activity score matrix of the snATAC-seq modality and Y_jk_ denote the gene expression matrix of the Stereo-seq modality. The average expression value for each gene k in the two modalities is defined as:

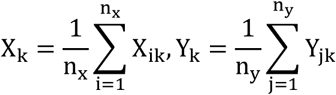

where n_x_ and n_y_ represent the number of cells in the snATAC-seq and Stereo-seq modalities, respectively.

The average capture rate, p̅ (p_mean_), is defined as the ratio of the average gene expression values between the two modalities and calculated as:

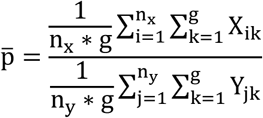

Based on this, the gene-specific perturbation factor r_k_ is defined as:

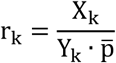

The cell-specific average capture rate p_i_ is calculated by matching the i-th quantiles of the library sizes between the query modality (ln_RNA,j_) and the reference modality (ln_ATAC,i_). This ratio is used to adjust the gene expression distribution of the reference modality. The formula is as follows::

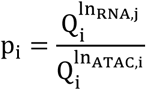

The cell-specific perturbation factor r_i_ is obtained by dividing by p_mean_:

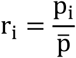

Finally, by combining the cell-specific perturbation factor and the gene-specific perturbation factor, the gene expression matrix is adjusted to generate the gene-cell perturbation ratio:

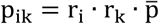

Sampling of the original snATAC-seq data is performed through the perturbation matrix, simulating a distribution that is closer to the target modality.

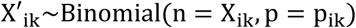

Data perturbed X′_ik_ in this manner serves as a reference, enhancing the model’s robustness to noise in real biological data and providing a complex scenario simulation for experimental design and biological hypothesis testing.

Next, we similarly trim the Stereo-seq data based on the set of highly variable genes derived from the reference modality, ensuring that its features are consistent with those of the snATAC-seq data for seamless cross-modal analysis. Additionally, the trimmed Stereo-seq data undergoes normalization and logarithmic transformation, using the same strategy applied to the snATAC-seq data.

#### Encoder and decoder

The autoencoder (AE) module consists of an encoder and a decoder, both utilizing fully connected layers to achieve dimensionality reduction and reconstruction of high-dimensional gene features. The encoder maps the input high-dimensional gene expression data (with dimensions corresponding to the number of genes) to a lower-dimensional latent space (with dimensions corresponding to the number of cell types) using a single fully connected layer, with the bias set to False. In the latent space, U_ic_ and V_jc_ represent cell-type-specific metagene expressions for snATAC-seq and Stereo-seq, which are weighted combinations of different genes, W_0_ denotes the weight matrix of the encoder as follows:

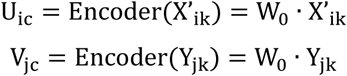

As the key component for dimensionality reduction, the encoder’s fully connected layer extracts cell-specific features related to cell types through linear mapping, while incorporating Dropout (dropout rate = 0.8) during the mapping process to enhance the model’s generalization ability and reduce the risk of overfitting.

The decoder employs a symmetric structure, with a fully connected layer that maps the latent variables from the low-dimensional metagene features back to the original high-dimensional space. The decoder’s task is to supervise the encoder by reconstructing the input reference modality gene expression matrix, ensuring that the latent variables extracted by the encoder preserve the major biological information.W_1_ denotes the weight matrix of the decoder as,

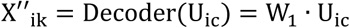

The use of fully connected layers not only simplifies the model design but also enhances the ability to adapt to the dimensionality reduction characteristics of different modalities, enabling the encoder and decoder to exhibit strong robustness in handling high-dimensional sparse data.

#### Discriminator

The design of the discriminator module is based on the low-dimensional metagene features generated by the encoder, serving as an integral part of the adversarial mechanism. Its task is to distinguish the modality source of the metagene features. The input to the discriminator is the latent variables generated by the encoder, and the output is the probability of modality labels. During adversarial training, the encoder attempts to generate modality-agnostic metagene features to "fool" the discriminator, making it difficult to accurately determine the modality source, thereby achieving cross-modal feature alignment in the latent variable space. W_2_ denotes the weight matrix of the discriminator:

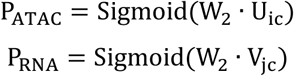

#### Loss function design

##### 1. Discriminator Loss

The discriminator loss is used to measure the discriminator’s ability to distinguish between two modalities. The goal is to maximize the probability of correct classification while ensuring that the contributions of different modalities are balanced. The formula for calculating the discriminator loss is as follows:

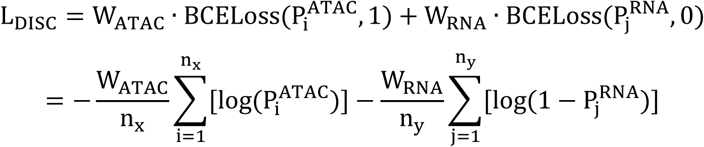

Where:

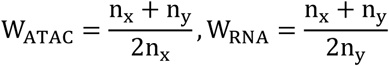

##### 2. Cross-Entropy Loss

This loss is used to evaluate the difference between the model’s predicted categories and the true categories. The formula for calculating the cross-entropy loss is as follows:

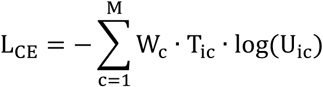

Where T_ic_ denotes the true label indicating whether the observed sample j belongs to category c. To address the class imbalance issue, we utilize class weights W_c_ to adjust the contribution of each class to the loss. The class weights are calculated as the inverse of the frequency n_c_ of each class:

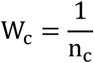

##### 3. Reconstruction Loss

The reconstruction loss is used to ensure that the output of the autoencoder is as close as possible to the input reference data, thereby preserving key information and minimizing information loss. The basic form of the reconstruction loss is the mean squared error (MSE), and its formula is as follows:

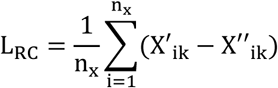

##### 4. Adversarial Loss

The adversarial loss is used to optimize the feature extractor in adversarial learning, making its generated features difficult for the discriminator to distinguish between different modalities. This is achieved by training the feature extractor to "fool" the discriminator, producing modality-agnostic features. The formula is as follows:

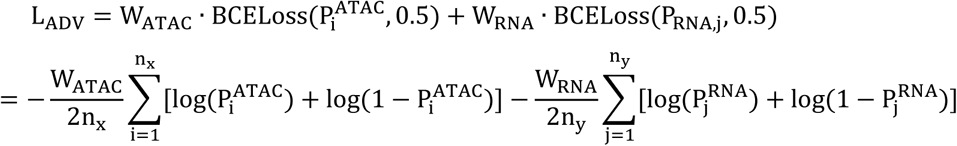

##### 5. Entropy Regularization Loss

Entropy regularization is used to encourage a more uniform distribution of the model’s output categories, thereby preventing overly confident predictions. The formula for calculating the entropy regularization loss is as follows:

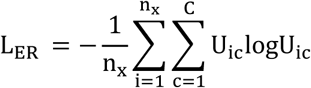

##### 6. L2 Regularization Loss

L2 regularization is used to constrain the magnitude of the model parameters, preventing overfitting. The formula is as follows:

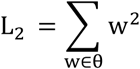

Where w represents the model parameters, which are the training parameters of the autoencoder.

#### Model Training

In the training process, a two-step training approach is employed, optimizing the discriminator and the feature extractor separately. This training strategy aims to achieve effective integration of the feature spaces from the two modalities (snATAC-seq and Stereo-seq) through adversarial learning, while simultaneously performing cell type classification and the reconstruction of input gene expression data. The training parameters set for the entire process include 1000 epochs, the Adam optimizer, and a learning rate of 0.01. Regularization parameters such as the L2 regularization coefficient are set to 0.01, and the dropout rate is set to 0.8.

In the first step, the parameters of the feature extractor (autoencoder module) are fixed, and only the discriminator is optimized. The task of the discriminator is to learn to distinguish the features of data from different modalities, with the input being the features extracted by the autoencoder from both modalities (e.g., snATAC-seq and Stereo-seq). The discriminator’s loss function is based on binary cross-entropy, aiming to maximize the accuracy of modality discrimination, while balancing the class imbalances caused by unequal sample sizes through a weighted loss. Through optimization, the discriminator gradually learns how to distinguish the features of the two modalities, providing a clear optimization target for the adversarial optimization of the feature extractor in the next step. The loss function for the first step is as follows:

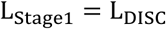

In the second step, the parameters of the discriminator are fixed, and only the feature extractor is optimized. The task of the feature extractor is to generate modality-agnostic shared features, while simultaneously performing cell type classification and high-fidelity reconstruction of the input data. To achieve this, the optimization of the feature extractor includes a weighted combination of multiple loss functions:

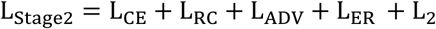

These two steps are carried out sequentially. In each training round, the discriminator is first updated to enhance its ability to distinguish between modalities, followed by the update of the feature extractor to make the modalities indistinguishable. After multiple rounds of alternative training, the model reaches a balance: the shared features generated by the feature extractor both preserve biological information and are modality-agnostic; the discriminator can no longer improve its classification accuracy, indicating that modality alignment has been achieved.

### One-to-one pairing between omics data

The metagene generated U_ic_,V_jc_ by the encoder is used for cell type prediction and cell matching. We employ the optimal transport algorithm for cross-modal mapping. The goal of optimal transport is to minimize the distance between cells. The key difference from traditional optimal transport algorithms lies in the computation of the cost matrix. Based on U_ic_, V_jc_, we rank the different cell types for each cell according to their probabilities, with the cell type showing the highest probability first and the lowest last, resulting in a type ranking vector sU_ic_,sV_jc_ for each cell. For two cells, *i* and *j*, from different modalities, if their top B_ij_ most likely cell types align, then the cost matrix is defined as:

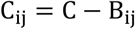

Next, to solve the optimal transport problem, the following objective function is constructed:

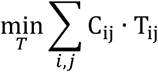

T_ij_ represents the transport amount from cell i to cell j. We use Earth Mover’s Distance (EMD) to solve this. For each cell j in Stereo-seq, we search for the cell i in snATAC-seq with the highest transport amount. In this way, each cell on the spatial transcriptome map was paired with chromatin accessibility from the snATAC-seq data. The chromatin accessibility of CREs was thus quantified in different layers and regions based on this mapping result.

### Layer and regional preference of various cell types and corresponding CREs

Based on one-to-one cell pairing result between snATAC-seq data and Stereo-seq data, we obtained the spatial distribution of 230 cell types among six cortical layers and 143 brain regions, classified into 9 cortical lobes, including prefrontal, frontal, cingulate, somatosensory, insular, auditory, temporal, parietal, and occipital lobes.

The cell density per cell cluster in six layers was calculated as the number of cells in the layer divided by the area size of the certain layer. The preference of each cell cluster in different brain regions was calculated as follows:

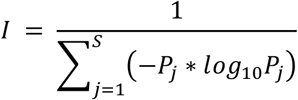

Here, *I* represented specificity index among cortical lobes, *P*_*j*_ is the proportion in a single brain region j, S is the number of brain regions the cell cluster was distributed.

### Cross-species analysis of various cell types

Human and mouse snATAC-seq data were retrieved from previously published studies^11, 15^ and downloaded from the chromatin accessibility database (http://catlas.org/humanbrain and http://catlas.org/mousebrain) and the GEO database^11^ (accession number: GSE246791). We applied the subsampling method to ensure similar numbers of nuclei across species at the subclass level based on previous studies^6^. Specifically, the species with the largest number of clusters under a given subclass was allowed a maximum of 200 nuclei per cluster. The remaining species then split this theoretical maximum (200 nuclei multiplied by the maximum number of clusters under the subclass) evenly across their clusters. The intersection of homologous genes among the three species according to the Ensemble ortholog database (version 91) were used for cross-species analysis, and gene activities of the 14,382 orthologous genes were quantified from chromatin accessibility of snATAC-seq data using “addGeneScoreMatrix” and “getMatrixFromArrow” function of ArchR. We then performed data normalization using “NormalizeData” function. Further integration analysis was conducted with Seurat “IntegrateData” function with top 2000 repeatedly variable genes as features. The proportion of nuclei overlap between species was calculated by the number of cells clustered in the same clusters of the integrative clustering result divided by the total number of cells for specific cell types from two species, using the "compare_cl" function from the published pipeline^6^. MetaNeighbor analysis was run to systematically assess the similarity between cell types within species. The data slot of integrated assay was reformatted and used as input with the “SummarizedExperiment” function of SummarizedExperiment (v.1.28.0) package. We calculated the AUROC score by the “MetaNeighborUS” function of MetaNeighbor (v.1.18.0) package with the parameter “one_vs_best=TRUE”.

### Cross-species analysis of CREs

The human and mouse CRE lists were download from http://catlas.org/humanbrain and http://catlas.org/renlab_downloads/wholemousebrain/sa2.subclassv3.final.peak.srt. First, we converted macaque and mouse CREs to human coordinates using the liftOver method with minMatch = 0.3 and UCSC chain files (“macFas5.hg38.all.chain”, “hg38.MacFas5.all.chain”, “mm10.hg38.all.chain”, and “hg38.mm10.all.chain”). Next, we reciprocally lifted the accessible elements back to raw genomes and only kept the regions that mapped to their original loci. We further removed converted regions with length > 1kb or in chrM, chr_random, chr_Un, and chr_alt chromosomes. Finally, we compared the homologous CREs using intervene^98^ (v.0.6.5) with f = 0.1. CREs showing both sequence similarity and open chromatin between two species were considered as conserved, whereas CREs with no homologous sequence or no open chromatin peaks in the other species were considered as species-specific. In this way, we classified CREs into 7 categories: Human/Macaque/Mouse-conserved (HMm, mammal-conserved), Human/Macaque-biased (HM, Human/Macaque-biased), Human/Mouse-specific (Hm), Macaque/Mouse-specific (Mm), Human-specific, Macaque-specific, Mouse-specific. We then calculated the proportion of these 7 categories of CREs among all CREs in each cell subclass.

In addition, we download the known transposable elements annotation file from UCSC. Only 4 TE types were kept for further analysis, including "DNA", "SINE", "LINE", "LTR". We calculated the percentage of CREs located in TE in these 7 categories, and performed pairwise Wilcoxon rank sum test with "bonferroni" correction for multiple testing among various categories.

### Comparison of LAMP5 and LAMP5/LHX6 between species

Human and mouse snATAC-seq data were retrieved from previously published studies^11, 15^, and we filtered to retain cells originating from the cortex. Despite some differences in annotation between datasets, we identified the most comparable subclass groups using gene activity scores of classical markers while preserving the original labels from the previous studies for visualization.

In order to compare the cell composition of LAMP5 and LAMP5/LHX6, we merged these two subclasses to calculate their proportions. To facilitate downstream computations, 5,000 cells were randomly sampled from the merged data for each species. Gene activity scores were normalized using a log2 transformation, and we used Seurat and Signac R package to perform integration.

To identify the primate-enriched motifs, we utilized Jaspar motifs (JASPAR2020, all vertebrate) to generate a motif matrix and motif object that was added to the Seurat object using Signac “AddMotifs” function. For motif activity scores, chromVAR was carried out according to default parameters with the Signac “RunChromVAR” function on the respective peak count matrices. The motif matrices of human, macaque, and mouse were merged, and differential enrichment was identified using the “FindMarkers” function with the mean.fxn parameter set to rowMeans.

### Characteristic of SST layer and GRN construction

We re-clustered the SST cell subclass as described above to identify cell groups with layer preferences. The previously published study had identified the different layer-specific markers of SST neurons^21^, so we examined the gene activity score of these layer markers and assign different cell types to the respective layers.

The GRN (gene regulatory network) construction was performed as described above. Briefly, we used CellOracle to calculate the transcription factor (TF)-target gene correlations in different SST groups with layer preferences. For visualization, we ranked the TF by their “eigenvector_centralit” and selected the top 5 TFs. At the same time, we also ranked the target genes by “coef_abs” and selected the top 5 genes. In the network, the target genes which were identified as layer markers in the previous study^21^ was highlighted.

### Correlation of OPC-specific and OLG-specific CREs with cortical hierarchy levels

Hierarchy levels of cortical regions in the visual system were derived from Felleman and Van Essen^51^. Based on one-to-one cell pairing result between snATAC-seq data and Stereo-seq data, we extracted the chromatin accessibility matrix of OPC-specific CREs in OPC cells and OLG-specific CREs in OLG cells. Then, we calculated the average chromatin accessibility for each OPC-specific CRE among all OPC cells and OLG-specific CRE among all OLG cells in various cortical regions. We applied the Spearman analysis to calculate the correlation between averaged chromatin accessibility and the hierarchy level for each CREs using the “cor.test” function. The CREs showing positive or negative correlation with the hierarchy levels were identified with cutoffs of *P* value < 0.01, R > 0.4 or R < -0.4. GO enrichment analysis was performed using these CREs associated genes by clusterProfiler (v.4.6.0).

### LDSC cell type specific analysis

To enable comparison with GWAS data of human phenotypes, we utilized the ‘liftOver’ with default parameters to convert cell type-specific CREs from “Macaca_fascicularis_5.0” to hg19 genomic coordinates via a chain file “macFas5ToHg19.over.chain” download from UCSC. We further filtered out converted regions exceeding 1 kb and retained only coordinates on each standard chromosome (chromosomes 1-22).

We used Linkage Disequilibrium Score Regression Analysis (LDSC) to assess the extent of genetic polymorphism enrichment associated with chromatin accessible regions differing among various cell types and traits. Summary statistics for GWAS of brain disorder, neurological traits, control traits were obtained from the GWAS Catalog and publications (Supplementary Table 7). We prepared the summary statistics in the standard format for linkage disequilibrium score regression and downloaded the relevant genotype resource from the pipline vignettes at https://github.com/bulik/ldsc. Using genotype data from the “1000G_Phase3_plinkfiles.tgz” and haplotype information from ‘hapmap3_snps.tgz’ as reference panels, we generated binary annotations for the homologous DNA sequences of each cell type and estimated LD scores on each standard chromosome. For each trait, we used the baseline model “1000G_Phase3_baseline_ldscores.tgz” along with the standardized regression weights “weights_hm3_no_hla.tgz” to partition phenotypic heritability into cell type-specific annotations, using the superset of all candidate regulatory peaks as background controls.

### Variant to CRE mapping

To broadly map the genomic features of disease risk signals across species, we first extracted all potential pathogenic variants from the GWAS summary data that passed the statistical test (P-value ≤ 0.05). Next, we used the R package “SNPlocs.Hsapiens.dbSNP144.GRCh37” along with their rsid information to retrieve the coordinates of these signals in the hg19 reference genome from the "dbSNP144" database. Subsequently, we performed intersection analysis between these coordinates and Cis-Regulatory Elements (CREs) in the Macaca fascicularis genome to identify disease risk-related CREs that may have similar functions across different species.

### Gene set related to brain disorder

The gene set related to brain disorder was obtained from Disease Ontology (DO). In R, we used the getHgDisease function from the geneset package to load the DO database by specifying source = "do". We then queried the DOID (Supplementary Table 7) and employed the transId method from the genekitr package, setting transTo = "symbol" and unique = TRUE, to retrieve the gene symbols.

### Sequence evolutionary conservation analysis

The phyloP score is a conservation score that quantifies the evolutionary conservation of each nucleotide by comparing the aligned sequences across multiple species^58, 59^. By evaluating phyloP scores, we assessed the evolutionary conservation of CREs associated with disease risk loci and those not associated with disease, in the context of cell-type genetic identity across species, relative to a shuffle background.

To focus the nucleotide evolutionary conservation assessment on comparisons between the Macaca fascicularis and human and mouse genomes, we first downloaded whole-genome masked sequences for three species (macFas5, mm10, and hg38) from UCSC, retaining the standard chromosome assemblies (sequences prefixed with "chr") for alignment analysis. Next, we performed reference-based alignment using the ‘LASTZ’ software, with parameter settings according to UCSC guidelines. For the closely related species macFas5 and hg38, we used the parameters “M=254 K=3000 L=3000 Y=3400 E=30 H=2000 O=400 T=2”; for the more distantly related macFas5 and mm10, we used the parameters “M=0 K=3000 L=3000 Y=9400 E=30 H=2000 O=400 T=1”. Subsequently, we applied ‘MULTIZ’ to assemble the pairwise ‘LASTZ’ alignment results into a multi-genome alignment output and set the phylogenetic tree for the three species to “((macfas5, hg38), mm10)” using the query from https://lifemap-ncbi.univ-lyon1.fr.

For evolutionary model generation, we used ‘phyloFit’ with the parameters ‘-i MAF --tree "((macfas5, hg38), mm10)" --subst-mod REV’; then, we used ‘phyloP’ to generate per-base phyloP score data, with the parameters ‘--mode CONACC --method LRT --wig-scores’ to compare the evolutionary conservation of the macaque genome against human and mouse genomes. Finally, we obtained the average phyloP scores for all CREs using ‘bigWigAverageOverBed’.

In the statistical analysis, for each cell type’s diff-CREs that co-localized with disease risk signals, we calculated its average phyloP score. To ensure a fair comparison of the evolutionary conservation of these Risk CREs, we performed 1000 bootstrap resampling on other diff-elements not belonging to Risk CREs and the supercluster of peaks, each time sampling an equal number of elements as the Risk CREs. We then aggregated the average scores from the 1000 re-samplings for statistical analysis.

### Gene Ontology (GO) enrichment

Gene Ontology annotation of interested CREs was performed using R package rGREAT^99^ (v.2.2.0). We downloaded AH111915 (Macaca fascicularis) database as reference using AnnotationHub (v. 3.8.0). For the pathway gene sets of Gene Ontology database, we extended the genes from its transcription start site with default parameters of "basal_upstream = 5000" and "basal_downstream = 1000". The total CREs of corresponding cell clusters were used as the background peak set for each. GREAT analysis was performed with "great" function of "min_gene_set_size = 5".

### RNA Scope

Macaque #3 was transcardially perfused with diethyl pyrocarbonate-treated PBS (DEPC-PBS), and then followed by 4% DEPC-PFA. The brain was removed and immediately embedded in optimum cutting temperature compound (OCT, SAKURA, cat#4583). The brain samples were stored at -80 °C. During samples processing, the OCT blocks were sectioned at 15 μm using a RWD freezing microtome, and brain sections were fixed onto adhesion microscope slides. The slides were baked at 60°C overnight. Then, RNA Scope Pretreatment Kit (Advanced Cell Diagnostics, cat#322381 and cat#322000) was used to unmask target RNA and permeabilize cells. Subsequently, RNA Scope Target Probe are hybridized to target RNA molecules. RNA Scope Detection Reagents (ACD, cat#322310) amplify the hybridization signals via sequential hybridization of amplifiers and color label probes. The RNA Scope procedure was performed according to the manufacturer’s instructions (https://acdbio.com). The details about the probes employed in the experiments are as follows: Probes against CPA6 (ACD, cat#1843691-C1) and MEF2C (ACD, cat#1843711-C2) were used to identify V1-specific L4 neurons in macaque monkeys; Probes against DCC (ACD, cat#1843701-C1) were used to identify layer4 neurons in macaque monkeys;

### Data visualization

For bigwig files, presented by Signac package "BigwigTrack" function. For the putative gene regulatory networks, presented with Cytoscape (v.3.10.1) tools. The residual graphs were plotted with R package ggplot2 (v.3.4.1) and pheatmap (v.1.0.12).

## Data availability

The processed data ready for exploration could be accessed and downloaded via https://macaque.digital-brain.cn/chromatin-accessibility (reviewers can download the data using the following ids and passwords: All raw data have been deposited to CNGB Nucleotide Sequence Archive (accession code: CNP0005530) and are publicly available as of the date of publication.

## Code availability

All data were analyzed with standard programs and packages, as detailed above, and the corresponding codes were accessible through https://github.com/JuneMeng/snATAC_MacaqueCTX_Atlas. Custom code of cell-cell pairing of snATAC-seq and spatial transcriptome data is available at https://github.com/sunyk740/RMGE/.

Additional information required to reanalyze the data reported in this paper is available from the lead contact upon request.

## Acknowledgements

The project was supported by National Science and Technology Innovation 2030 Major Program (STI2030-2021ZD0200100), National Natural Science Foundation of China (NSFC) Outstanding Youth Foundation (No. T2422026), National Key R&D Program of China No.2022YEF0203200, National Natural Science Foundation of China (No. U23A6010, 32100487) and Shanghai Science and Technology Development Funds (23QA1410400).

## Author contributions

Methodology, J.M., C.C., Z.Z., Y.S., K.H. and T.F.; Investigation, J.M., Y.L., C.C., Z.Z., Y.B.; Formal Analysis, J.M., C.C., Z.Z., Y.S. and Y.H.; Writing – Original Draft, Y.L. and Y-D.S.; Writing – Review & Editing, J.M., T.F., Z.L., Z.S., L.L., C.L., and M.P.; Conceptualization and supervision, S.L. and Y-D.S..

---

**Extended Data Fig. 1.**
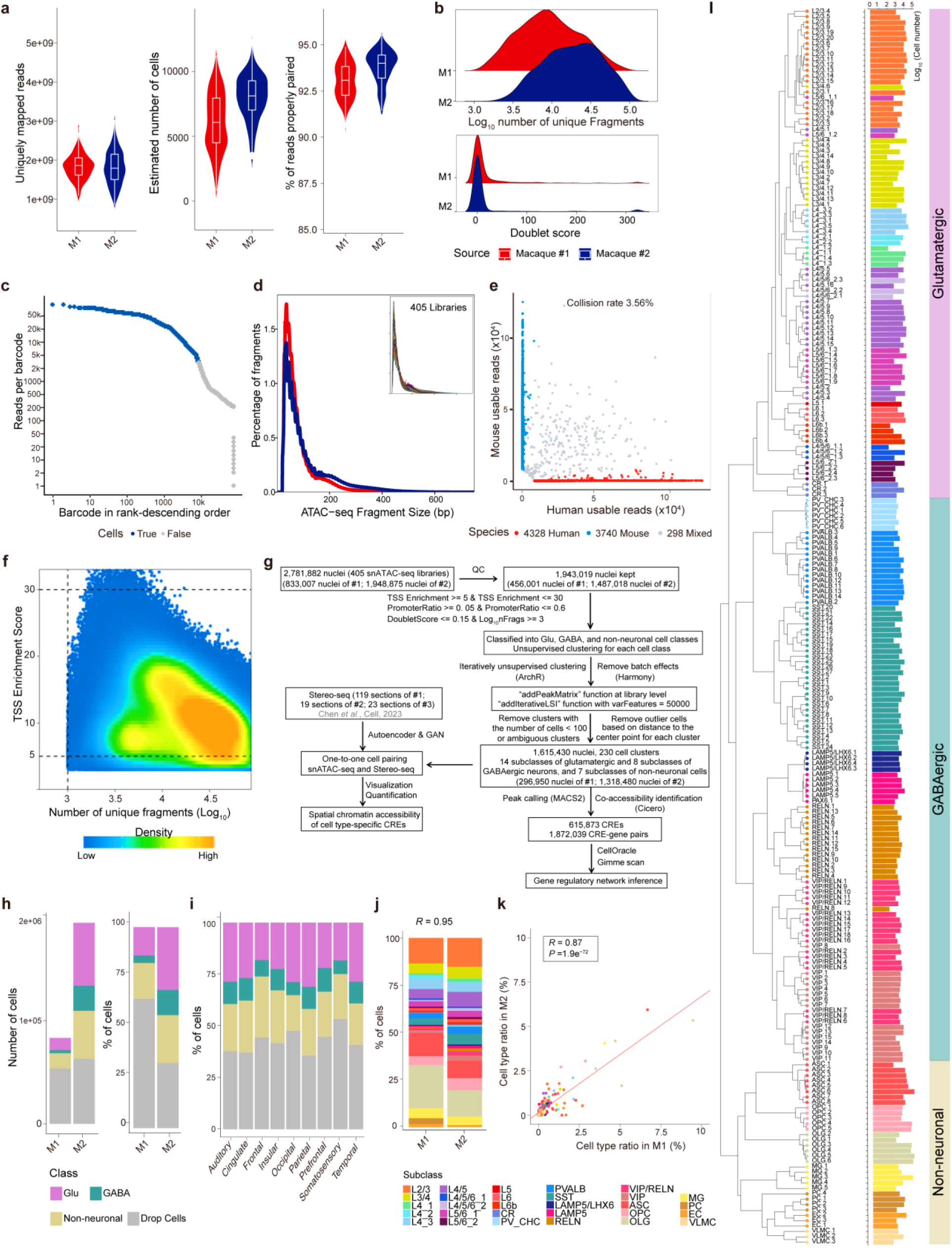
Quality control metrics of snATAC-seq data. **a**, Box plots showing the distribution of uniquely mapped reads (left), estimated numbers of cells (middle) and percent of reads properly paired (right) in biological replicates Macaque #1 and #2 of snATAC-seq libraries from each tissue sample. **b**, Density plots showing the distribution of number of unique fragments (upper) and doublet score (lower) in biological replicates Macaque #1 and #2 of snATAC-seq libraries. **c**, Knee plot for the detection of cells by the relationship between reads per barcode vs. the number of barcodes. **d**, Frequency distribution plot showing the fragment size distribution of each snATAC-seq library. **e**, Fraction of cell collision estimated from species mix samples of human and mouse. **f**, Density plot illustrating fragments per nucleus and individual TSS enrichment. Nuclei in middle right quadrant were selected for analysis (5 < TSS enrichment < 30 and number of unique fragments > 1000). **g**, The quality control process for the snATAC-seq dataset. **h**, Bar plot showing the numbers (left) and percentages (right) of glutamatergic, GABAergic and non-neuronal cells in biological replicates Macaque #1 and #2. **i**, Bar plot showing percentages of glutamatergic, GABAergic and non-neuronal cells in 9 cortical lobes. **j**, Bar plot showing percentages of various cell clusters in biological replicates Macaque #1 and #2. **k**, Dot plot showing the relationship of cell type compositions between Macaque #1 and #2. Correlation coefficient and *P* value was correlated by Pearson correlation test. **l**, Hierarchical clustering of 230 cell subclusters based on chromatin accessibility and the number of nuclei in each subcluster.

**Extended Data Fig. 2.**
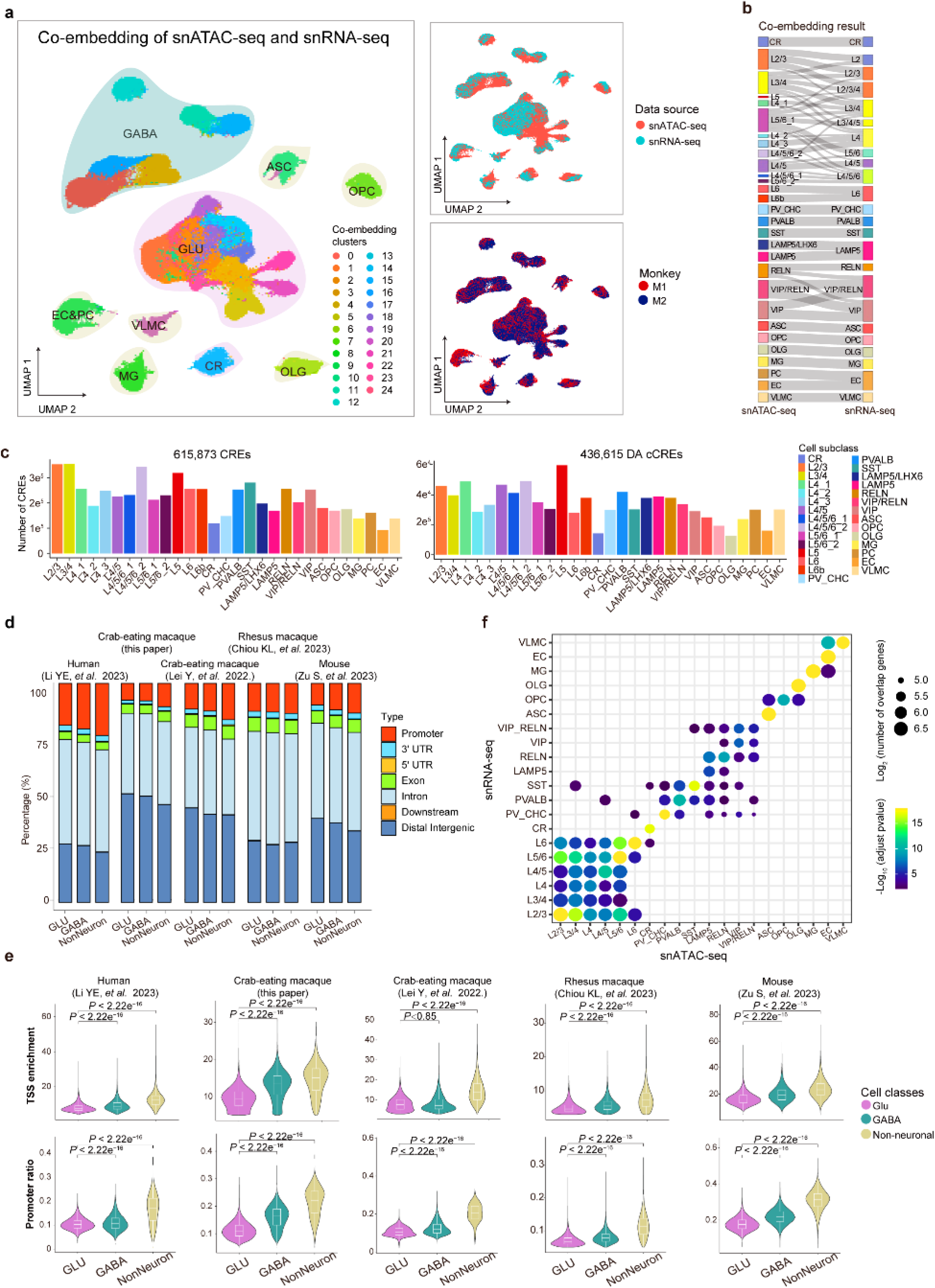
Statistics of CREs across various cell types in the macaque cortex. **a**, UMAP showing the co-embedding of snATAC-seq and snRNA-seq data from the entire macaque cortex (left), and UMAP layout showing single nuclei from different cortical lobes (top right) and different monkeys (bottom right). **b**, Correspondence of snATAC-seq-revealed cell clusters with snRNA-seq-identified cell types. **c**, The number of identified CREs and differential accessible peaks in various cell subclasses. **d**, Bar plots showing the fraction of CREs that overlaps with different classes of annotated sequences in glutamatergic, GABAergic and non-neuronal cells in our macaque datasets and published snATAC-seq datasets of human, macaque and mouse brain^11, 15, 18, 19^. **e**, Comparison of the TSS enrichment and promoter ratio of glutamatergic, GABAergic and non-neuronal cells in our macaque datasets and published snATAC-seq datasets of human, macaque and mouse brain^11, 15, 18, 19^. **f**, Dot plotshowing the overlap of distal CREs linked genes and differentially expressed genes between snATAC-seq and snRNA-seq identified cell types. *P* values were calculated by hypergeometric test.

**Extended Data Fig. 3.**
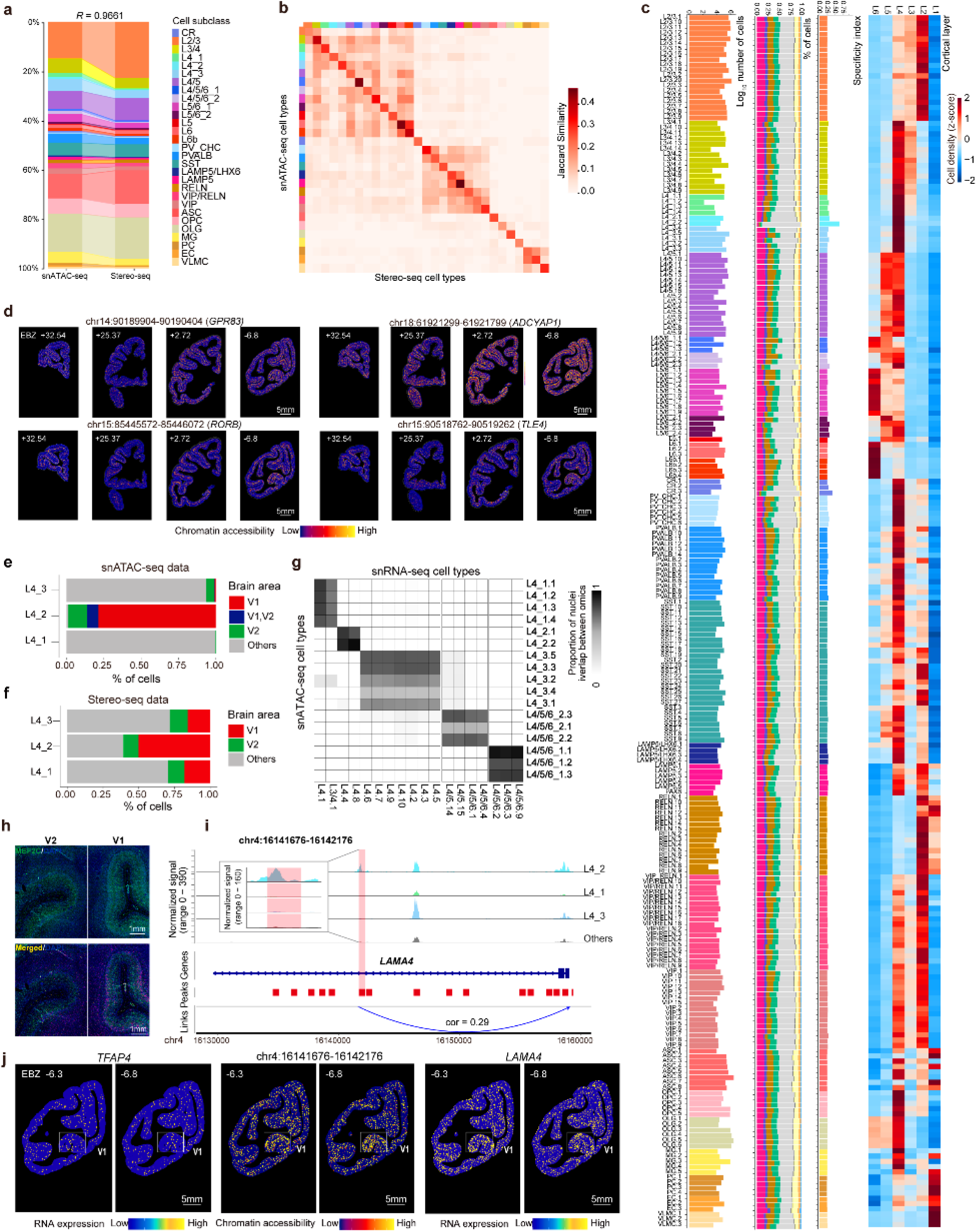
Spatial mapping of chromatin accessibility of CREs in various cell types. **a**, Correspondence of percentages of various cell clusters revealed by snATAC-seq clustering and spatial mapping result on the Stereo-seq map. **b**, Heatmap showing the Jaccard similarity of marker genes identified from snATAC-seq data and spatial transcriptome data for the same cell cluster. **c**, Layer- and region-distribution of snATAC-seq-defined cell clusters by spatial mapping. The first to fourth columns from left to right represent the number of cells in each cluster, the regional distribution preference, percentage of cells in nine cortical lobes and the relative density of cells in six cortical layers for each cell type. **d**, Spatial maps showing the chromatin accessibility of four example CREs shown in Fig. 2e across cortical layers in representative coronal sections at indicated EBZ coordinates. **e**, Bar plots showing the relative distribution of three L4 clusters across cortical brain regions in the snATAC-seq dataset. **f**, Bar plots showing the relative distribution of three L4 clusters across cortical brain regions in the spatial transcriptome mapping result. **g**, Heatmap showing the correspondence of snATAC-seq-identified cell clusters and transcriptome-defined cell types. **h**, RNAscope experiment results showing the expression pattern of the *MEF2C* gene in V1 and V2 regions of the macaque cortex. Probes against *MEF2C* (ACD, cat#1843711-C2) was list in Supplementary Table 8. DAPI was used for nuclear counterstaining. **i**, Genome browser track view showing a differential accessible distal CRE linked to *LAMA4* across three L4 glutamatergic clusters and other glutamatergic clusters. **j**, Spatial maps of expression patterns of *TFAP4* and *LAMA4*, and chromatin accessibility of the CRE linked to *LAMA4* at two representative sections (EBZ -6.3 and -6.8).

**Extended Data Fig. 4.**
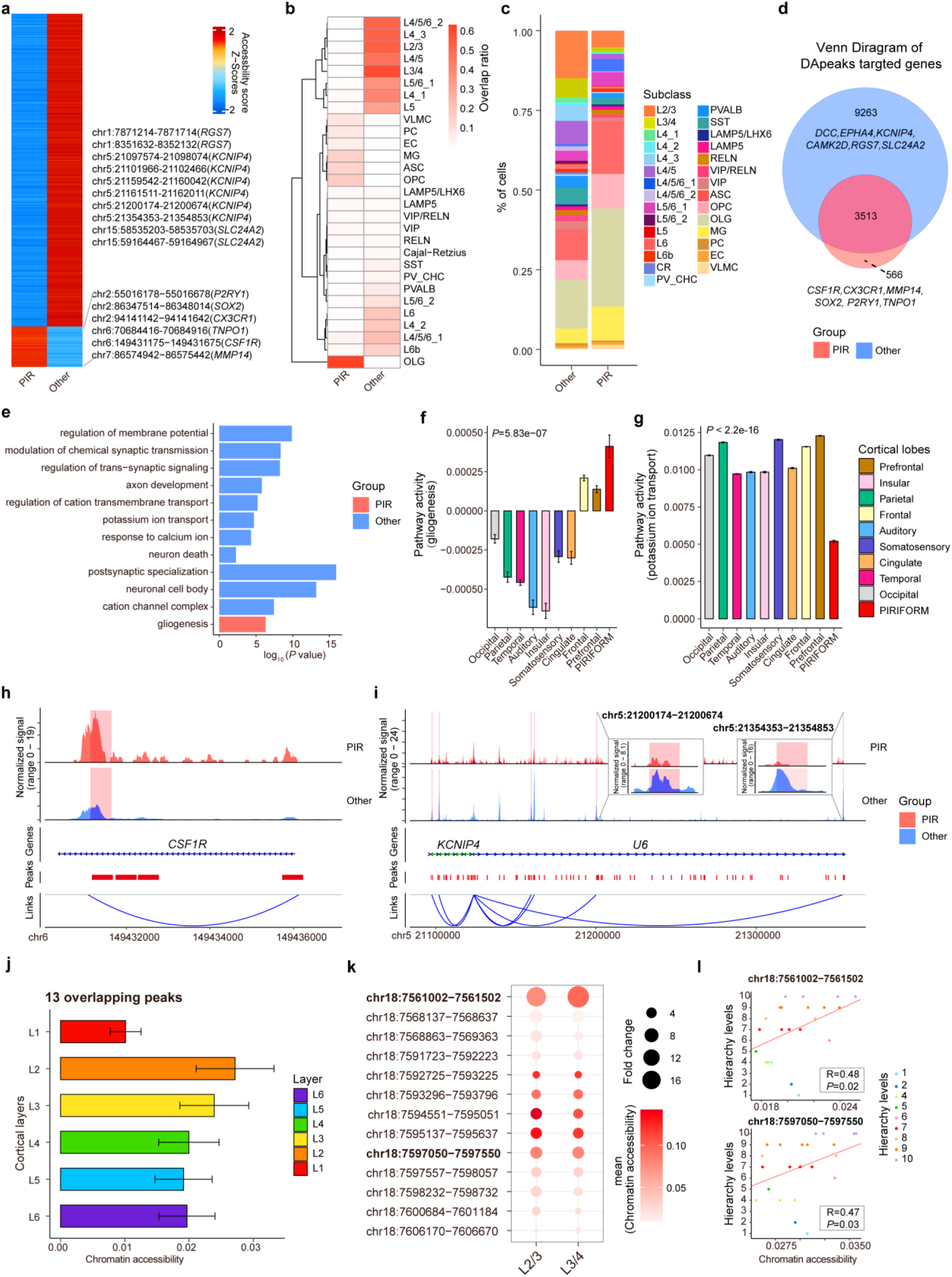
Characterization of CREs for the piriform cortex and *CBLN2 gene*. **a**, Heatmap showing the chromatin accessibility of differential CREs between piriform cortex and all the other neocortex regions. **b**, Heatmap showing the overlap between the up-regulated CREs in piriform cortex and the other neocortex regions and the differentially accessible CREs in various cell types. **c**, Bar plot showing percentages of various cell clusters in piriform cortex and the other neocortex regions. **d**, Venn plot showing the overlap of genes linked by up-regulated CREs in piriform cortex and the other neocortex regions. **e**, Pathway enrichment for genes linked to the up-regulated CREs in piriform cortex and the other neocortex regions shown in **d**. **f-g**, Barplot showing the pathway activity of “gliogenesis” (**f**) and “potassium ion transport” (**g**) across nine cortical lobes. The pathway activity was calculated by averaged expression level of enriched genes in the pathway subtracted by the aggregated expression of control feature sets. **h**, Genome browser tracks showing one up-regulated CRE in piriform cortex linked to the TSS region of *CSF1R*. **i**, Genome browser tracks showing the down-regulated CREs in piriform cortex linked to the TSS region of *KCNIP4*. **j**, Bar plots showing the quantitative chromatin accessibility level of the 13 overlapping peaks in our macaque dataset with the CREs of *CBLN2* in *Shibata M, et al. Nature. 2021* across six cortical layers. Error bars represent standard error of the mean (SEM). **k**, Dot plot showing the the fold change in the relative accessibility of L2/3, L3/4 subclass versus other cells. Color indicates the mean accessibility score. Size indicates the fold change. All comparisions are statistical significant (Wilcoxon rank-sum test). **l**, Dot plots showing the correlation of chromatin accessibility and the hierarchy levels of various brain regions in the somatosensory system.

**Extended Data Fig. 5.**
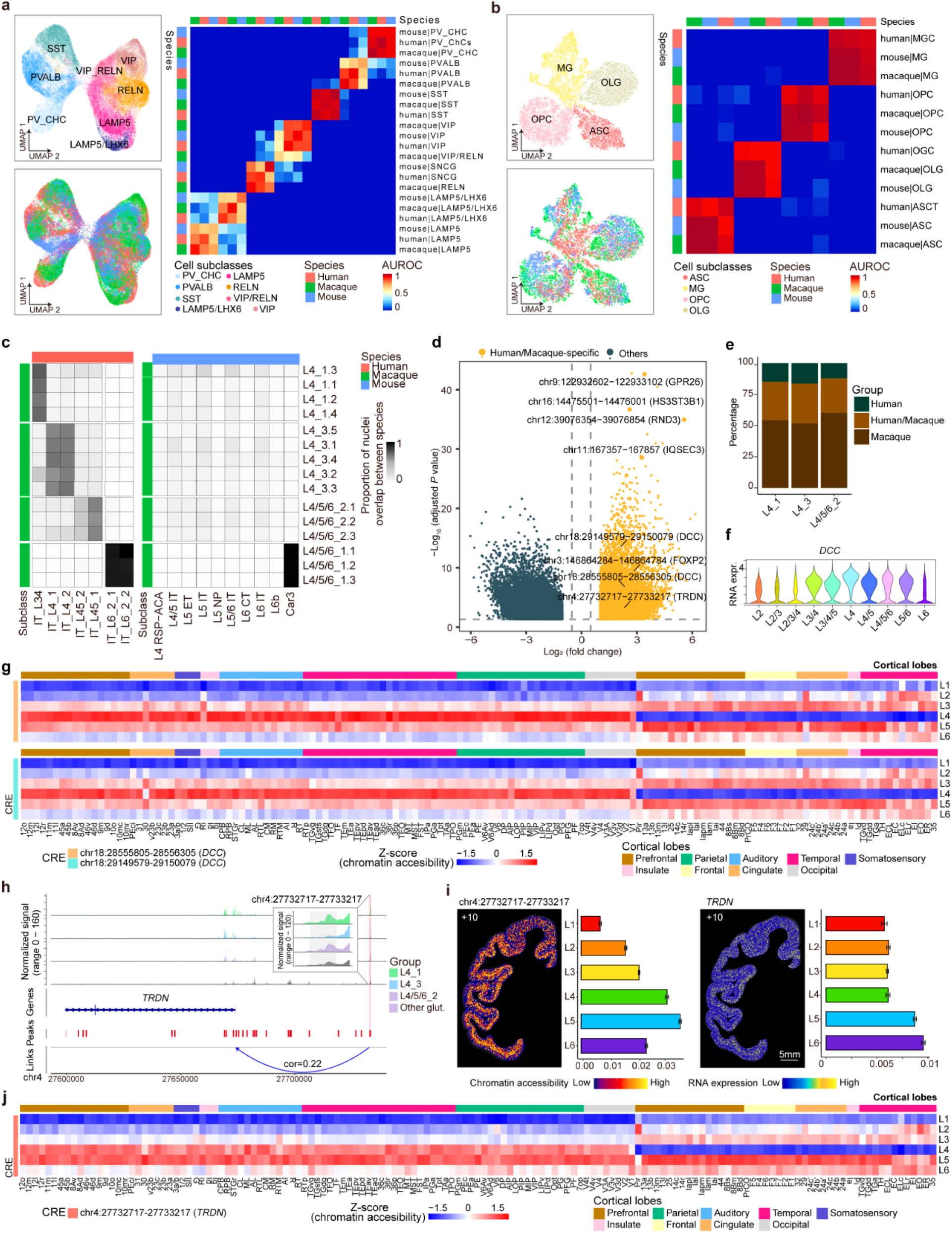
Cross-species comparison of CREs for glutamatergic neurons. **a**, **Left**, UMAP co-embedding of GABAergic neurons (upper) and non-neuronal cells (below) from snATAC-seq data of macaques, humans and mice. Single nuclei were colored by their cell subclasses. **Right,** Heatmap showing the similarity of snATAC-seq derived GABAergic cell clusters across macaques, humans and mice using MetaNeighbor. **b**, Similar to (**a**), but for non-neuron cells. **c**, Heatmap depicting the pairwise comparisons of snATAC-seq derived glutamatergic cell types between macaques and humans or mice. **d**, Volcano plot showing the differentially accessible CREs between human/macaque-biased and other snATAC-seq-defined cell types. **e**, Bar plots showing the conservation between humans and macaques for CREs with high chromatin accessibility in human/macaque-biased cell types shown in **d**. **f**, Violin plots showing the expression level of *DCC* among various glutamatergic neuron subclasses of the macaque snRNA-seq data. **g**, Heatmap showing the average chromatin accessibility of CREs associated with *DCC* across cortical layers and regions. **h**, Genome browser tracks showing the chromatin accessibility of human/macaque-biased CRE linked to *TDRN* in the human/macaque-biased and other cell types in the macaque cortex. **i**, Spatial map showing the chromatin accessibility of the human/macaque-biased CRE shown in **g** and the expression level of *TDRN* in a representative Stereo-seq section at indicated coordinate. The right panel showing the quantitative chromatin accessibility or gene expression level across six cortical layers. **j**, Heatmap showing the average chromatin accessibility of CRE linked to *TRDN* across cortical layers and regions.

**Extended Data Fig. 6.**
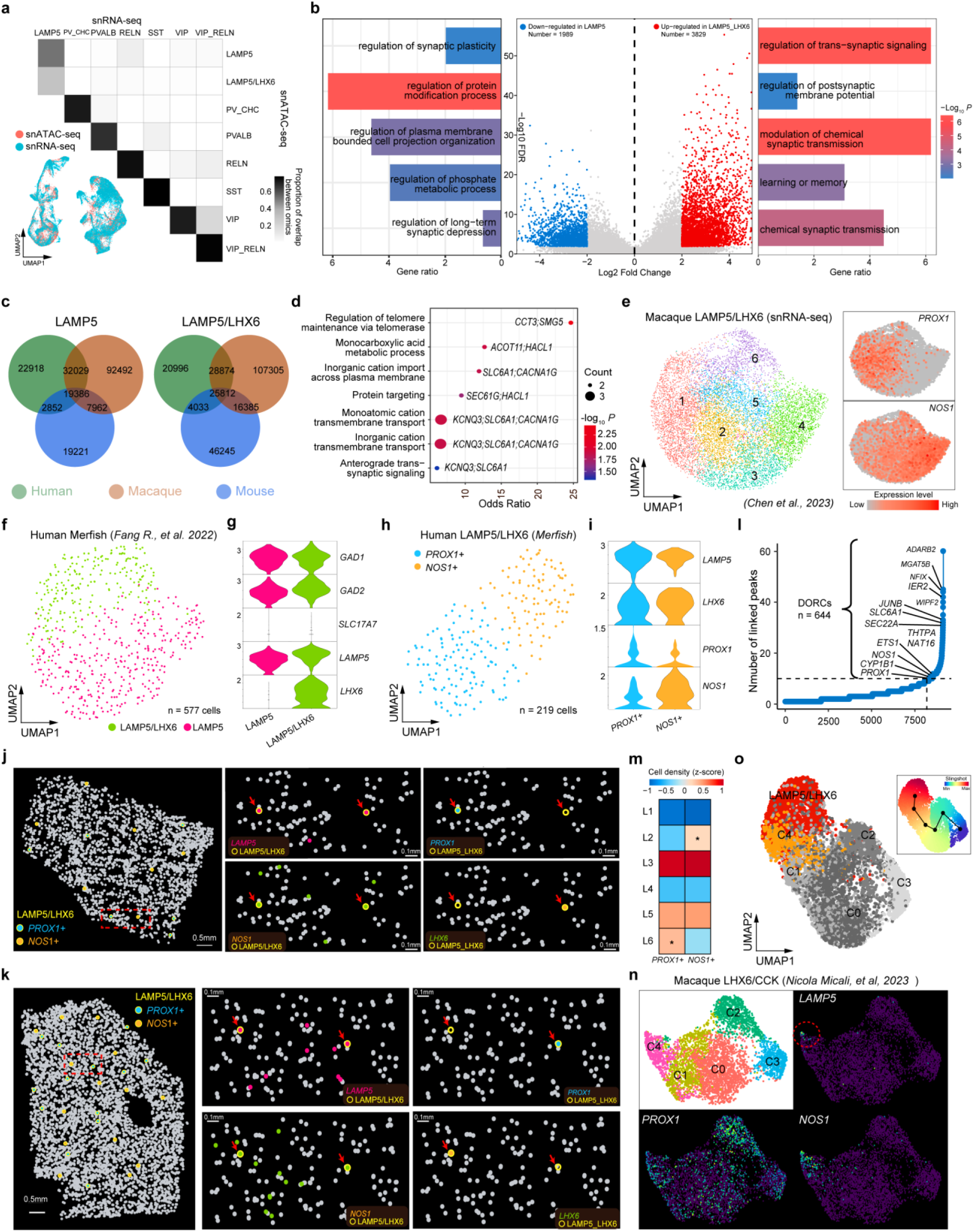
Characterization of CREs in LAMP5/LHX6 GABAergic neuron subgroups. **a,** UMAP embedding of snATAC-seq and snRNA-seq GABAergic cells from macaque cortex (lower left). Heatmap of overlap scores showing the correspondence between snRNA-seq and snATAC-seq derived macaque GABAergic cell subclasses (upper right). **b**, Volcano plot showing differential accessible peaks between LAMP5 and LAMP5/LHX6 neurons (middle). GO pathway enrichment analysis of genes linked to distal DA peaks in LAMP5 (left bar plot) and in LAMP5/LHX6 (right bar plot). **c,** Venn plot showing the overlap of CREs among human, macaque and mouse LAMP5 (left), and LAMP5/LHX6 cells (right). **d,** Gene ontology enrichment of the shared CREs between human and macaque LAMP5/LHX6 neurons. **e,** UMAP embedding of snRNA-seq LAMP5/LHX6 neurons (left) and normalized gene expression of PROX1 and NOS1 (right). **f,** UMAP showing the LAMP5 and LAMP5/LHX6 cell clusters derived from MERFISH data of the human dorsolateral prefrontal cortex (dlPFC) (*Fang R., et al. Science. 2022*). **g,** Violin plots depicting the expression levels of marker genes for LAMP5 and LAMP5/LHX6 cell types. **h,** UMAP illustrating the PROX1+ and NOS1+ subpopulations within the LAMP5/LHX6 cell type (left), alongside their corresponding gene expression patterns (right). **i,** Violin plots depicting the expression levels of marker genes for PROX1+ and NOS1+ subclusters. **j-k**, Dot plots showing the spatial pattern of LAMP5/LHX6 cell types (left) and gene expression patterns for indicated genes (right) in the MERFISH tissue sections. The four insets on the right are magnified views of the region outlined by the red dashed box on the left, depicting the spatial expression patterns of the genes *LAMP5*, *LHX6*, *PROX1*, and *NOS1*, respectively. LAMP5/LHX6 cells were marked by yellow circles and red arrows. **l,** The number of significantly correlated peaks for each gene in LAMP5/LHX6 neurons. **m,** Layer distribution of the two distinct groups of LAMP5/LHX6. An asterisk (*) indicates a significant difference between the two groups in the given layer (*P* < 0.01). **n,** Re-clustering of MGE-derived LHX6/CCK neurons from development macaque brain identified five distinct subclusters (C0 to C4). Cluster C4 showed co-expression of *PROX1* and *NOS1* with relatively low levels of *LAMP5*. **o**, Integration of developmental clusters with adult LAMP5/LHX6 populations demonstrated preferential alignment of C4 with adult LAMP5/LHX6 neurons. The upper right panel shows the pseudotime trajectory inferred by Slingshot, ranging from early (blue) to late (red) stages.

**Extended Data Fig. 7.**
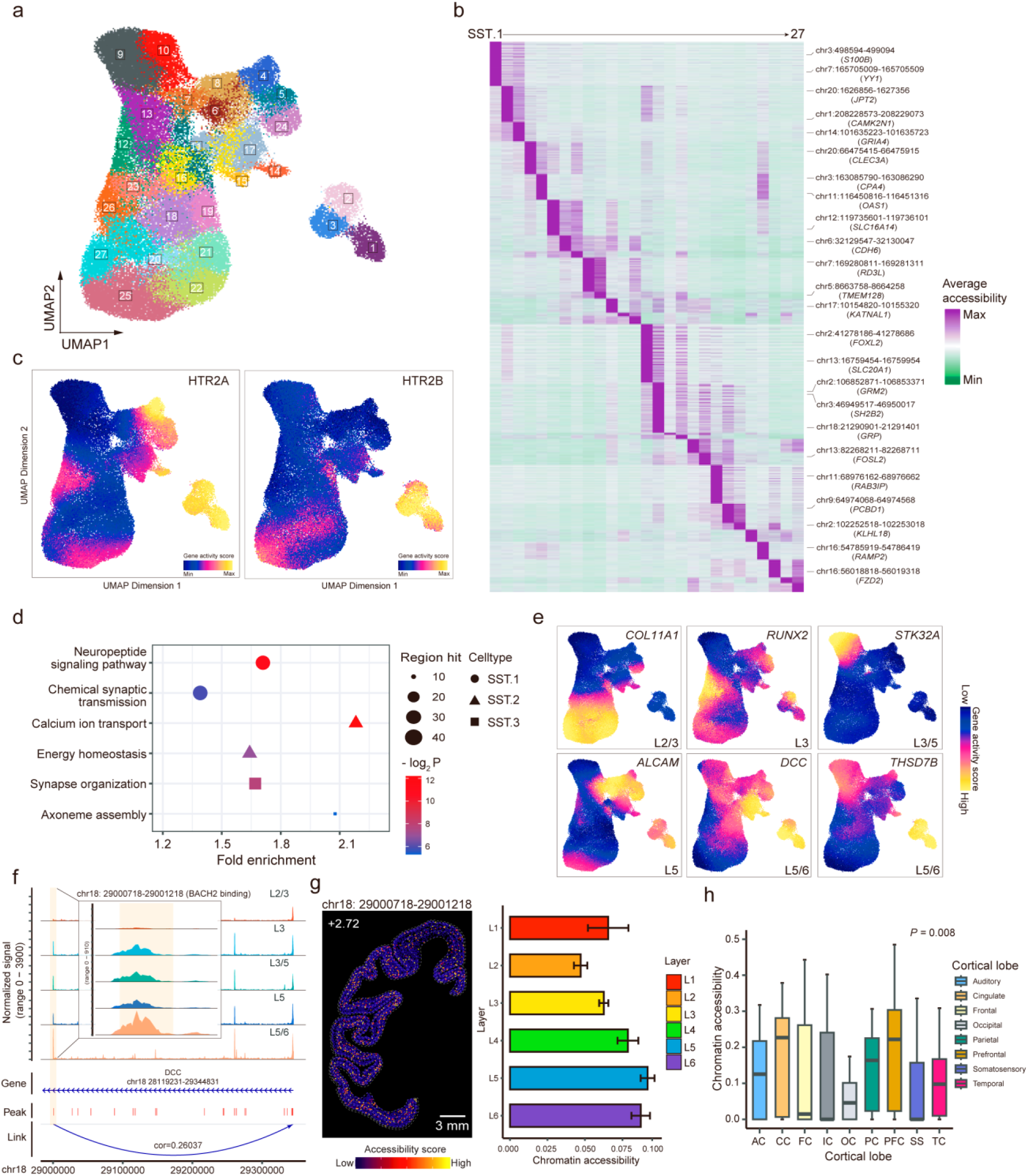
Characterization of CREs in layer-specific SST GABAergic neurons. **a,** UMAP embedding of snATAC-seq-based SST neurons. **b,** Heatmap of top differentially accessible (DA) peaks across 27 SST subclusters (1,000 cells randomly sampled per cluster) and distal linked genes are annotated. **c,** Normalized gene activity scores of *HTR2A* and *HTR2B* across SST subclusters. **d,** Representative Gene Ontology terms enriched by rGREAT analysis of DA peaks. **e,** Normalized gene activity scores of layer markers for SST neurons (right). **f,** Genome browser track view showing a differential accessible distal CRE linked to *DCC* across various SST cell types. **g,** Spatial visualization of chromatin accessibility for the CRE linked to *DCC* (shown in **j**) in representative section (left), and bar plots showing the average chromatin accessibility in six cortical layers for the CRE (right). **h,** Boxplot showing chromatin accessibility of the CRE associated with *DCC* (shown in **j**) in SST neurons across cortical lobes. *P*-values: Kruskal-Wallis test.

**Extended Data Fig. 8.**
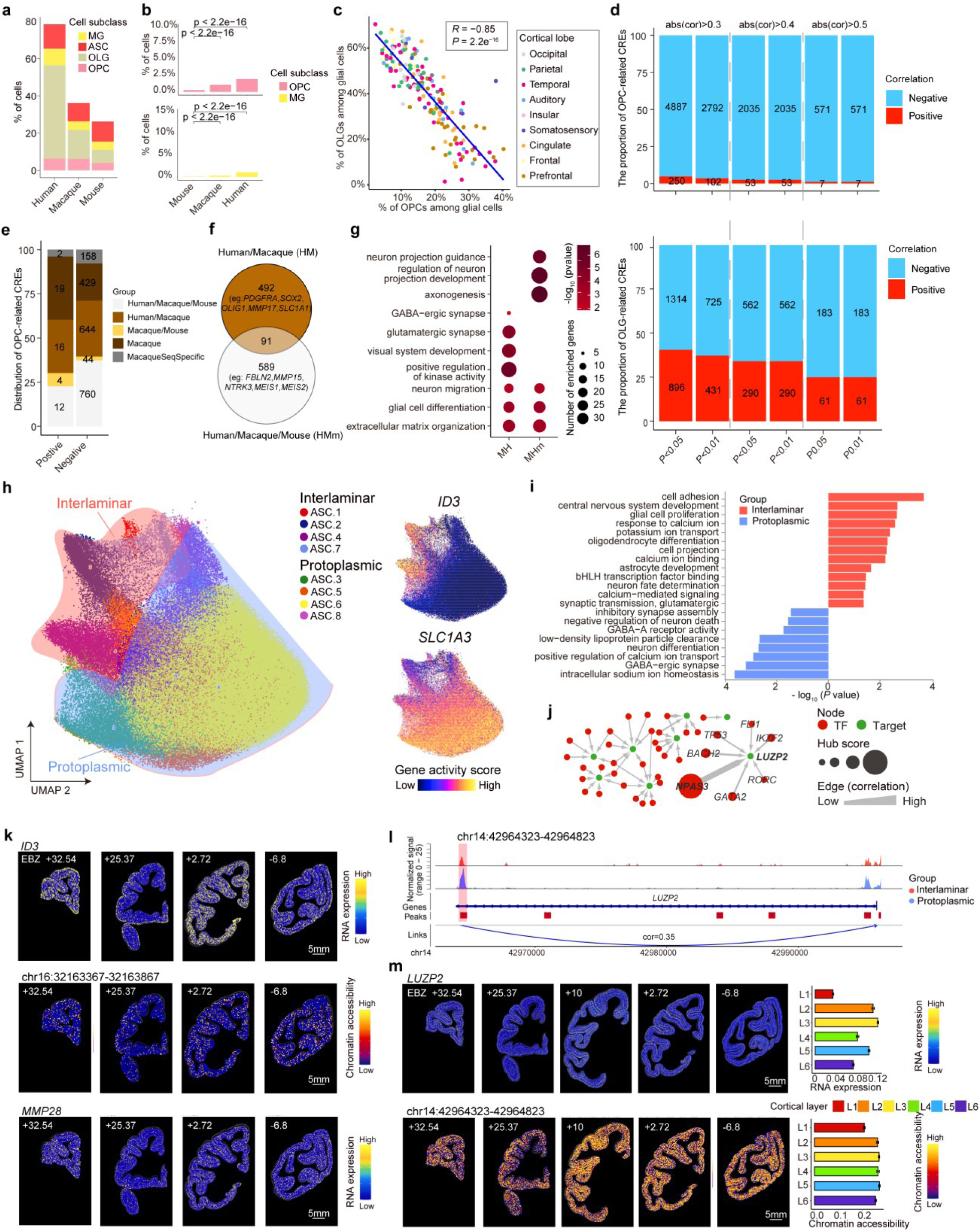
Characterization of CREs for cortical non-neuronal cells. **a**, Histogram showing the percentages of different glial cell subclasses based on snATAC-seq data across humans, macaques and mice. **b**, The percentages of OPCs and MGs based on snRNA-seq data across humans, macaques and mice. *P* values were calculated by Z-test. **c**, Dot plot showing the relative percentages of OPCs and OLGs among various macaque cortical regions using the snATAC-seq data. Coefficient and *P* value were calculated by Spearman correlation test. **d**, Bar plots showing the numbers and proportions of CREs in OPCs (upper) and OLGs (lower) with significantly negative or positive correlation (Spearman correlation) with the hierarchy level of cortical regions in the visual system. **e**, Bar plots showing the distribution in species-specific or mammal-conserved groups for CREs in OPCs negatively or positively (|*R| >* 0.4 and *P* < 0.01) correlated with the hierarchy level of cortical regions in the visual system. **f**, Venn diagram showing the relationship between human/macaque-biased and mammal-conserved CREs in OPCs negatively (*R* < -0.4 and *P* < 0.01) correlated with the hierarchy level of cortical regions in the visual system. **g**, Pathway enrichment of genes linked with human/macaque-biased and mammal-conserved CREs in OPCs positively or negatively correlated with the hierarchy levels of brain regions in the visual system. **h**, UMAP embedding of astrocytes (left) and normalized gene activity score of *ID3* and *SLC1A3* (right). **i**, Pathway enrichment of CREs upregulated in the interlaminar and protoplasmic ASCs. **j,** Gene regulatory network of TFs and targeted genes in protoplasmic astrocytes. **k**, Spatial maps of expression patterns of *ID3* and *MMP28*, and chromatin accessibility of the CRE linked to *MMP28* at four representative sections. **l**, Genome browser tracks showing the chromatin accessibility of CRE linked to *LUZP2* between protoplasmic and interlaminar astrocytes. **m**, Spatial maps (left) and quantification (right) of expression pattern of *LUZP2* and chromatin accessibility of the CRE linked to *LUZP2* at five representative sections. Error bars represent SEM.

**Extended Data Fig. 9.**
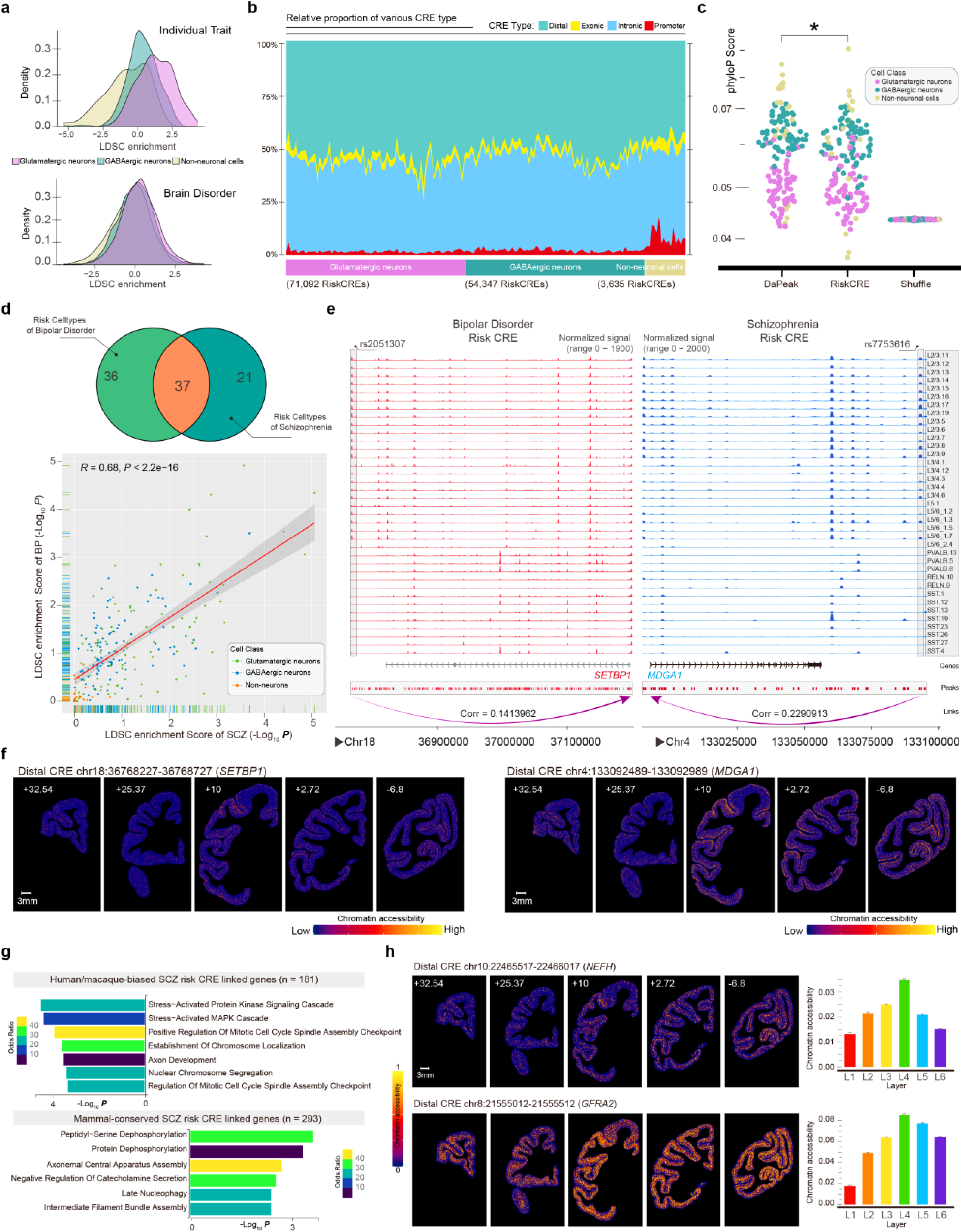
Cell type enrichment of CREs colocalized with disease-associated risk variants. **a**, The distribution of LDSC enrichment scores across glutamatergic neurons, GABAergic neurons and non-neuron for individual trait (upper) and brain disorders (bottom). **b**, The composition of brain disorder-associated CREs. **c**, Comparison of phyloP scores among Risk CREs (DaPeaks with disease risk SNPs), DaPeaks without disease risk SNPs, and a shuffled background. Significant differences were observed between Risk CREs and the DaPeaks (\**P* < 0.05; Wilcoxon rank-sum test). **d**, Venn plot showing the overlap between SCZ and BP risk variants enriched cells types (upper) and Pearson correlation between LDSC enrichment score of SCZ and BP (bottom). **e**, Genome browser tracks showing chromatin accessibility for gene *NT5DC1* and *SEMA6A*, CREs colocalized with risk variants of bipolar (left) and schizophrenia (right) in neuron cell types were highlighted. **f**, Spatial visualization of cell type-enriched CREs that co-localized with disease-associated SNPs loci in representative sections at different coordinates in EBZ numbers. **g**, Pathways enriched by genes linking to SCZ risk SNP associated CREs in L4_3 glutamatergic neurons. **h**, Spatial visualization of *NEFH*- and *GFRA2*-linked CREs associated with AD SNPs in representative EBZ sections (left). The bar plot illustrates the mean chromatin accessibility of CREs across different layers (right). Error bars represent SEM.

**Extended Data Fig. 10.**
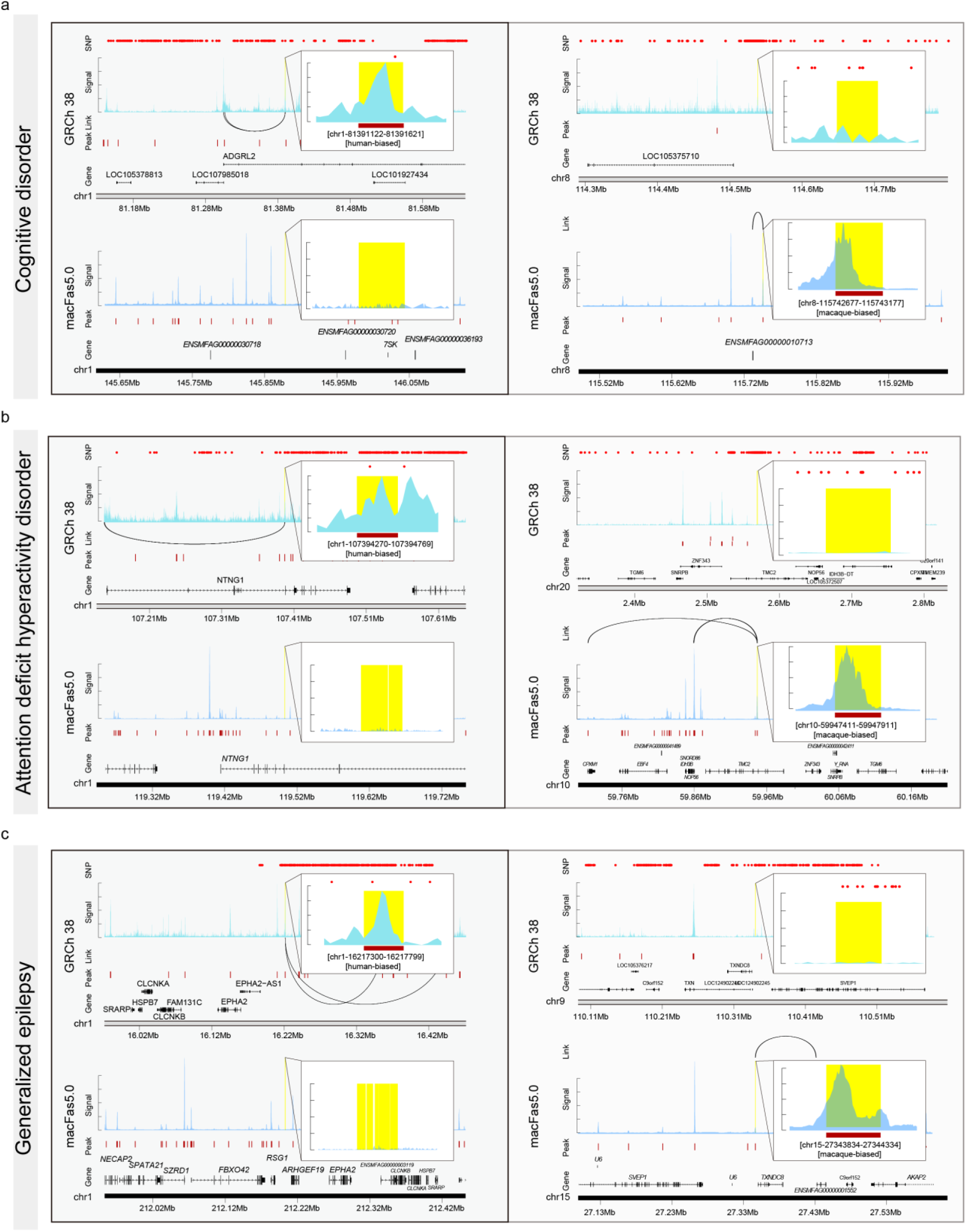
Genome browser tracks showing chromatin accessibility of human- and macaque-biased CREs co-localized with disease risk SNPs in LAMP5/LHX6 neurons. **a-c**, Genome browser tracks showing chromatin accessibility of human- and macaque-specific CREs in LAMP5/LHX6 neurons co-localized with risk SNPs of cognitive disorder (**a**), attention deficit hyperactivity disorder (**b**) and generalized epilepsy (**c**). Highlighted regions are indicated by color bands. **Left,** a human-active CRE (top) with a conserved but inactive/weakly active counterpart in macaque (bottom). **Right,** a macaque-active CRE (bottom) with a conserved but inactive/weakly active counterpart in human (top). Red dots at the top of each panel represent the genomic loci of risk SNPs.

## Supplemental tables

**Supplementary Table 1. Quality metrics of snATAC-seq libraries and brain region list.**

**Supplementary Table 2. Correspondence of snATAC-seq-identified cell clusters with snRNA-seq-defined cell types.**

**Supplementary Table 3. The cis-regulatory elements and their linked genes among various cell types.**

**Supplementary Table 4. Cross-species comparison of snATAC-seq data of humans, macaques and mice.**

**Supplementary Table 5. Putative gene regulatory networks across layer-specific SST neurons.**

**Supplementary Table 6. List of CREs in OPCs and OLGs positively and negatively correlated with hierarchy levels of various brain regions in the visual system.**

**Supplementary Table 7. The full list of linkage disequilibrium score regression analysis result.**

**Supplementary Table 8. Probes design for RNAScope Multiplex Assays.**

